# Deciphering the heterogeneity of differentiating hPSC-derived corneal limbal stem cells through single-cell RNA-sequencing

**DOI:** 10.1101/2023.11.24.568553

**Authors:** Meri Vattulainen, Jos G.A. Smits, Dulce Lima Cunha, Tanja Ilmarinen, Heli Skottman, Huiqing Zhou

**Affiliations:** Faculty of Medicine and Health Technology, Tampere University, Finland; Department of Molecular Developmental Biology, Faculty of Science, Radboud Institute for Molecular Life Sciences, Radboud University, The Netherlands; Department of Human Genetics, Radboud University Medical Center, The Netherlands

## Abstract

A comprehensive understanding of the human pluripotent stem cell (hPSC) differentiation process stands as a prerequisite for the development hPSC-based therapeutics. In this study, single-cell RNA-sequencing (scRNA-seq) was performed to decipher the heterogeneity during differentiation of three hPSC lines towards corneal limbal stem cells (LSCs). The scRNA-seq data revealed nine clusters encompassing the entire differentiation process, among which five followed the anticipated differentiation path of LSCs. The remaining four clusters were previously undescribed cell states that were annotated as either mesodermal-like or undifferentiated subpopulations, and their prevalence was hPSC line-dependent. Distinct cluster-specific marker genes identified in this study were confirmed by immunofluorescence analysis and employed to purify hPSC-derived LSCs, which effectively minimized the variation in the line-dependent differentiation efficiency. In summary, scRNA-seq offered molecular insights into the heterogeneity of hPSC-LSC differentiation, allowing a data-driven strategy for consistent and robust generation of LSCs, essential for future advancement toward clinical translation.

**Highlights:**

- hPSCs to LSCs spans epithelial, mesodermal, and undifferentiated cell states.
- scRNA-seq reveals the cell line-dependent differentiation heterogeneity.
- ITGA6 and AREG can be used to select pure LSC-like subpopulation.

## 1. Introduction

Corneal limbal stem cell (LSC)-driven repair mechanisms and homeostatic renewal of corneal epithelium (CE) are essential processes in the maintenance of human ocular surface health and clarity (Deng et al., 2019). Guiding human pluripotent stem cells (hPSCs) toward specific cell types such as LSCs possesses significant promise in translational applications and could be used for ocular surface regeneration for patients suffering from bilateral limbal stem cell deficiency (LSCD) (Ghareeb et al., 2020; Mahmood et al., 2022). Major advantages of hPSCs as the LSC source include abundance and donor-independent “off-the-shelf” nature, and to this day, multiple CE/LSC differentiation methods for hPSCs have been introduced (Mahmood et al., 2022). In an attempt to facilitate their clinical translation, some protocols have progressed from traditional reliance on conditioned media, feeder cells and/or serum (e.g. (Ahmad et al., 2007; Cieślar-Pobuda et al., 2016; Hayashi et al., 2012) towards simplified feeder- and serum-free versions such as our own (Hongisto et al., 2017; Mikhailova et al., 2014).

Despite ongoing improvements in our hPSC differentiation protocol and in those developed by others, there are persistent, universal challenges that continue to impede the effective implementation of hPSC-based therapeutics. Specifically, poor cost-efficiency arising from high variation between individual hPSC lines (Bock et al., 2011; Burrows et al., 2016; Kilpinen et al., 2017; Osafune et al., 2008), clones (D’Antonio et al., 2018) and differentiation batches (Volpato et al., 2018), coupled with variable differentiation efficiency towards the desired cell type will easily render promising protocols infeasible for widespread use. Additionally, residual cellular heterogeneity within hPSC-derived grafts may impair efficacy and most importantly, it poses a significant risk to patient health in terms of tumorigenic potential (Sato et al., 2019; Yamanaka, 2020).

State-of-the-art omics technologies such as single-cell RNA-sequencing (scRNA-seq) allow deciphering the heterogeneity of complex cellular systems and can serve as a powerful tool for assessing the efficiency of hPSC differentiation protocols (Cuomo et al., 2020). In recent years, the use of scRNA-seq techniques has significantly expanded our knowledge about the LSC identity and limbal progenitor distribution on the multistage corneal differentiation trajectory *in vivo* (e.g., Collin et al., 2021; Dou et al., 2021; Li et al., 2021; Ligocki et al., 2021). The corneal atlas generated from these works also forms a valuable up-to-date reference for the hPSC-derived LSCs (for a comprehensive recent review, see Arts et al., 2023).

Through years of dedicated work on hPSC-LSC differentiation in our laboratory, we have obtained promising results using traditional characterization methods such as immunofluorescence staining (IF) (Hongisto et al., 2017; Vattulainen et al., 2019, 2021). Although commonly known as a challenge, heterogeneity in the differentiation of hPSCs into LSCs has not been systematically addressed. For a comprehensive understanding of this matter, an unbiased deep molecular phenotyping approach is required. In this study, and for the first time, we used scRNA-seq to explore the heterogeneity of cell populations during hPSC-LSC differentiation and attempted to find feasible solutions for more consistent and robust derivation of hPSC-LSCs.

## 2. Results

### 2.1. Single-cell RNA-seq recapitulates the temporal marker gene expression patterns during the differentiation of hPSCs toward LSCs

In this study, we used one human embryonic stem cell (hESC) line Regea08/017 (hereafter, “hESC”) and two human induced pluripotent stem cell (hiPSC) lines WT001.TAU.bB2 and WT003.TAU.bC (hereafter, “hiPSC1” and “hiPSC2”, respectively) for differentiation toward LSCs with previously established protocol (Hongisto et al., 2017) (Figure 1A). The hESC line has been used for the initial establishment of the method (Hongisto et al., 2017; Mikhailova et al., 2014) and since then, has served as our golden standard line with consistently good performance in various experimental settings (Kauppila et al., 2023; Koivusalo et al., 2019; Sorkio et al., 2018; Vattulainen et al., 2019, 2021). The hiPSC1 line has been used for hPSC-LSC differentiation in one previous study (Vattulainen et al., 2021), while hiPSC2 is a newly established line (characterization described in Supplementary File 1). During differentiation, the morphological inconsistency between the cell lines was evident: by day 24 (D24), majority of hESC cultures consisted of epithelial monolayers with cuboid cell shape, whereas hiPSC1 and hiPSC2 yielded heterogeneous cultures with atypical PSC-like colonial growth (Supplementary Figure S1).

**Figure 1.**
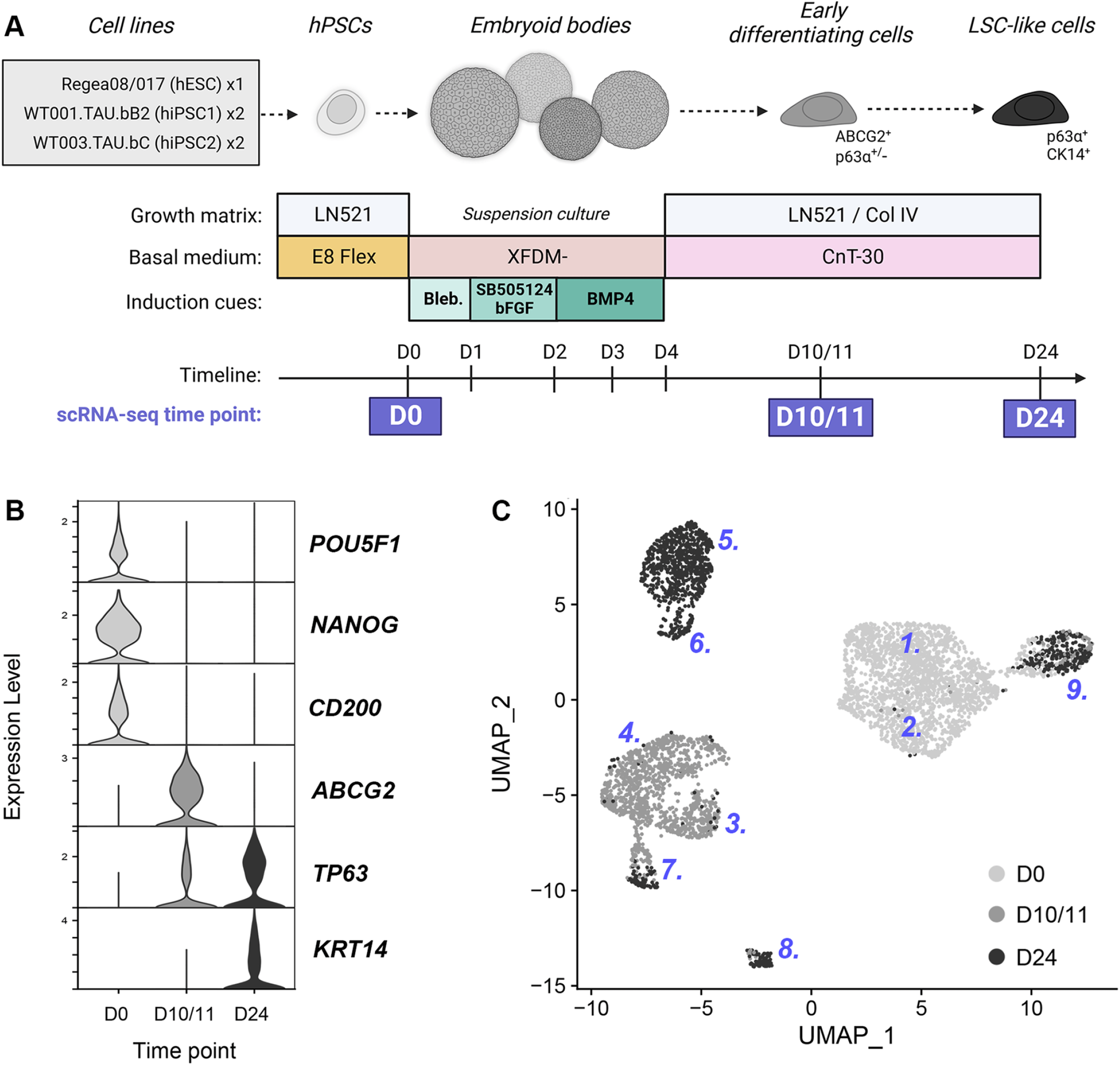
Single-cell RNA-sequencing (scRNA-seq) of three human pluripotent stem cell (hPSC) lines differentiated towards corneal limbal stem cells (LSCs). A) Schematic overview of the differentiation process, with scRNA-seq sample collection time points indicated at days 0, 10/11 and 24. Illustration created with Biorender.com. B) Expression of selected pluripotency- and LSC-associated marker genes in cells collected at the studied time points shown by violin plots. C) UMAP dimensionality reduction graph of the scRNA-seq data showing nine identified clusters.

To define the precise cell states within heterogenous populations, we carried out scRNA-seq for five cell batches, comprising one replica of the hESC and two replicas of both hiPSCs. Samples were collected from undifferentiated cells at day 0 (D0) and from two previously identified key time points at D10/11 and D24, as based on our previous observations these time points contain populations with distinct LSC-associated marker expressions (Vattulainen et al., 2019, 2021). After quality control (QC) and filtering of raw scRNA-seq data, 4840 high-quality cells were acquired for downstream analysis (Supplementary Figure S2). The scRNA-seq data confirmed the expected overall expression patterns for well-known marker genes over time (Figure 1B): decreased expression of pluripotency markers after D0, expression of previously detected LSC marker *ABCG2* (ATP-binding cassette subfamily G member 2) mainly at D10/11, and strong induction of LSC/progenitor marker *TP63* and *KRT14* (cytokeratin 14) expression at D24. Based on the gene expression profiles of the cells, a total of nine clusters were identified using Louvain clustering (Supplementary Figure S3) (Stuart et al., 2019) and visualized in high dimensional space using Uniform Manifold Approximations and Projections (UMAPs) (Figure 1C).

### 2.2. Single-cell RNA-seq elaborates the differentiation heterogeneity during the hPSC-LSC differentiation process

The nine clusters were subsequently characterized in detail in terms of the contributing time points, cell lines and their associated cell cycle phases (Figure 2A-B; Supplementary Figure S3; Supplementary Table S1). Six clusters comprised cells almost exclusively (>95%) from a single time point. The main difference between C1 and C2, collected at D0, arose from cell cycle and subtle cell line-specific variations (Supplementary Figure S3). Therefore, they were merged into a single cluster for downstream analyses (C1-2). Cluster 3 (C3) and Cluster 4 (C4) contained cells mostly collected at D10/11, with a few cells originating from D24, while Cluster 5 (C5) and Cluster 6 (C6) cells were collected solely at D24. Due to their association to certain collection days, we defined these six clusters (C1-2, C3, C4, C5 and C6) as “time point-specific”. Judging from the percentage of cycling cells in each time point specific cluster, the proliferation of the cells decreased over time. (Figure 2B, Supplementary Table S1)

**Figure 2.**
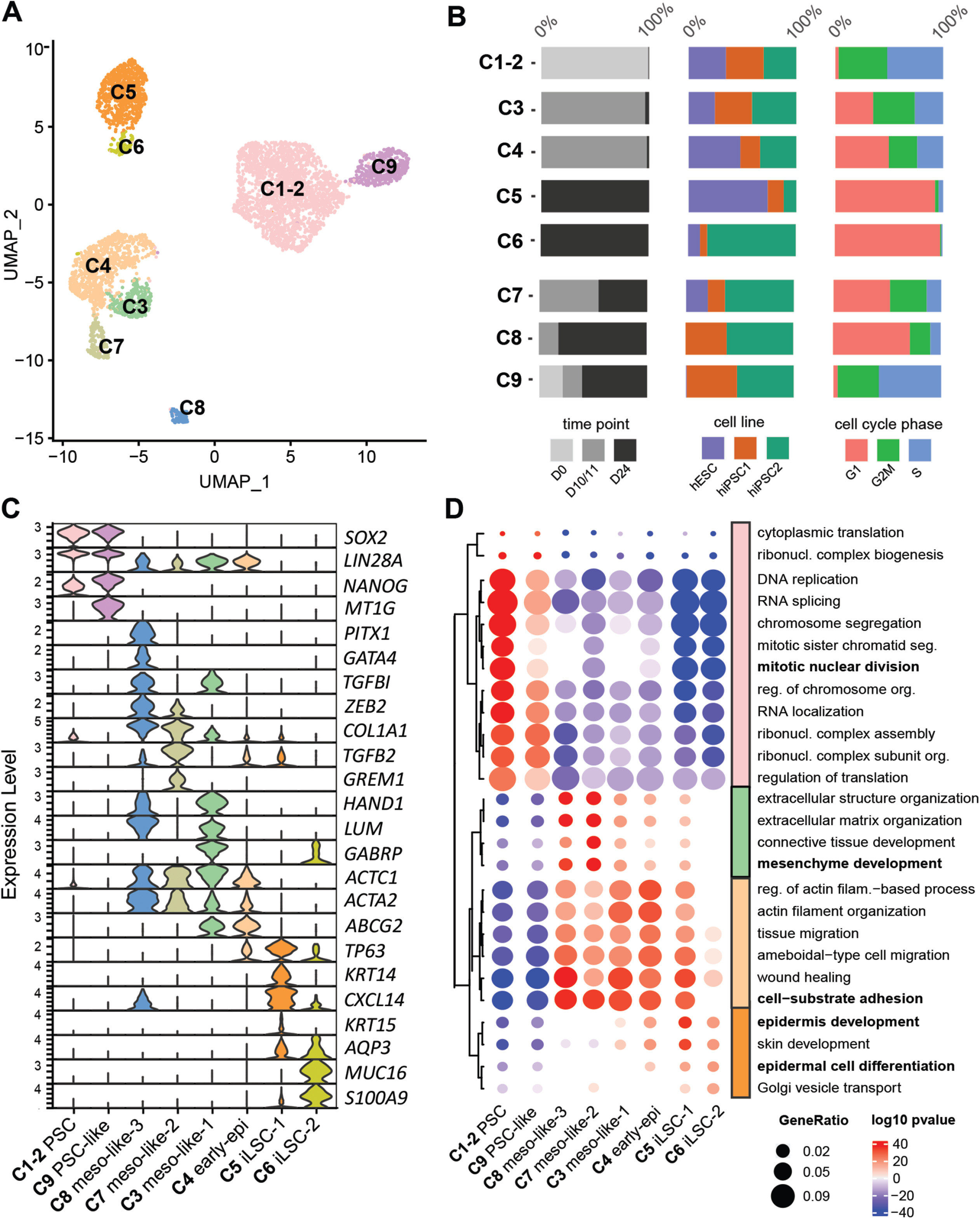
Single-cell RNA cluster characteristics and annotation. A) UMAP with cluster annotations. B) Distribution of cells in clusters per time point, cell line, and cell cycle phase. For details, see Table 1. C) Expression of selected marker genes across the clusters. D) Gene Ontology (GO) enrichment analysis based on positive cluster markers (red-pvall) and negative cluster markers (blue-pval). GO-terms are kmean clustered into 4 clusters.

Among the remaining three clusters, Cluster 7 (C7) and Cluster 8 (C8) contained a mixture of cells originating from both D10/11 and D24, while Cluster 9 (C9) comprised cells from all three time points. C7 contained slightly more cells from D10/11, whereas most cells from C8 and C9 were obtained at D24. Nevertheless, we defined this group as “mixed time point” clusters. There was a moderate number of cycling cells in C7 and C8, while C9 contained actively cycling cells at comparable levels with C1-2 cells. (Figure 2B; Supplementary Figure S3; Supplementary Table S1)

Importantly, the studied cell lines showed varying distributions to the identified clusters, with the largest overall contribution to the time point-specific clusters coming from hESC, and the majority of the cells in the mixed time point clusters derived from hiPSC2. First, the combined C1-2 contained similar number of cells from all three studied cell lines. In D10/11-associated clusters, however, hESC was slightly under-represented in C3 in comparison to the other two lines, but slightly over-represented in C4. Among the D24-associated clusters, C5 contained a clear majority of hESC-derived cells and rather small fractions from the two hiPSC lines. C6 was formed by a small total number of cells, which mainly originated from hiPSC2. Interestingly, C7 was the only mixed time point cluster containing a small fraction of hESC-derived cells, whereas both C8 and C9 were formed solely from hiPSC1 and hiPSC2. (Figure 2B; Supplementary Table S1)

### 2.3. Time point-specific clusters represent expected cell states involved in the LSC differentiation pathway

Next, we set out to identify the precise cell state in each cluster, using a combination of marker gene expression and Gene Ontology (GO) analysis (Gene Ontology Consortium, 2004). The protein-level expression of selected markers was subsequently confirmed by IF, and clusters were assigned with descriptive annotations.

Many C1-2 marker genes were associated with proliferation and there was high expression of pluripotency markers, including *SOX2* (SRY-box transcription factor 2) and *NANOG* (Figure 2C; Supplementary Figure S4). Marker genes were enriched for GO terms “mitotic nuclear division” and “DNA replication” (Figure 2D), which are functions associated with actively dividing cells, including embryonic stem cells. IF of undifferentiated hPSCs (at D0 time point), which constituted the C1-2, showed high-intensity, uniform expression of pluripotency-associated cluster markers TRA-1-81 (encoded by *PODXL*), OCT3/4 and LIN28 (Figure 3A; Supplementary Figure S4), consistent with the given annotation as “PSC”.

**Figure 3.**
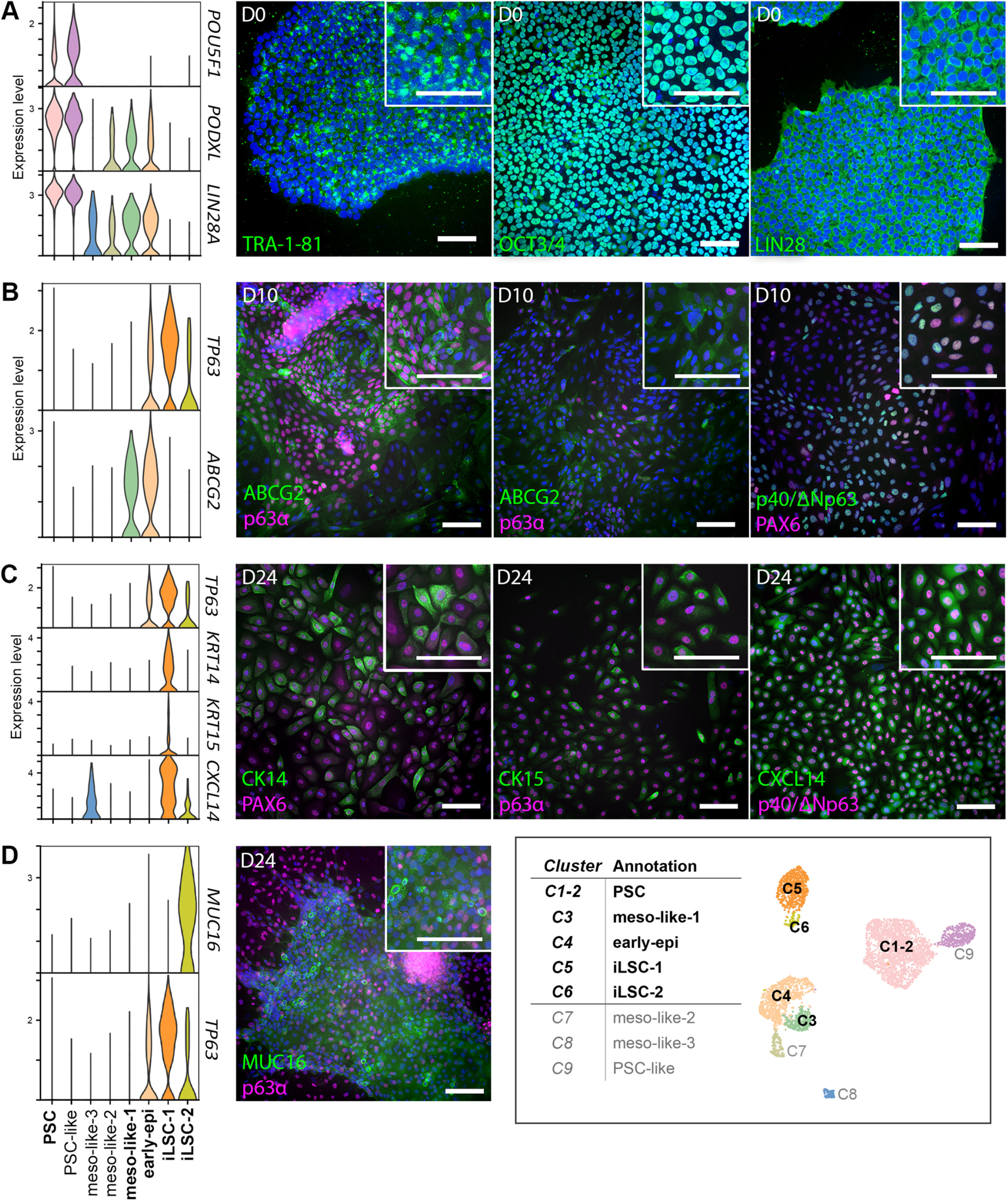
Immunofluorescence validation of time point-specific Clusters 1-6. A) Expression of pluripotency/Cluster 1/2 (PSC) markers *PODXL* (encoding TRA-1-81), *POU5F1* (encoding OCT3/4), and *LIN28A* (LIN28) in the undifferentiated hPSCs at D0 time point. B) Expression of Cluster 3 and 4 (early-epi and meso-like-1) marker *ABCG2* (ABCG2) and Cluster 3 marker *TP63* (encoding p63α and p40/ΔNp63) as well as corneal lineage marker *PAX6* (PAX6) at D10 time point. C) Expression of *PAX6* (PAX6) and Cluster 5 (iLSC-1) markers *KRT14* (encoding CK14), *KRT15* (encoding CK15), *TP63* (p40/ΔNp63), and *CXCL14* (CXCL14) at D24 time point. C) Expression of Cluster 6 (iLSC-2) marker *MUC16* (MUC16) at D24 time point. Representative IF images are shown for line hiPSC1. Cell nuclei counterstained with DAPI or Hoechst 333420 (both shown in blue). Scale bars, 100 µm in all images.

Both C3 and C4 had high expression of the stem cell marker *ABCG2* (Figure 2C; Supplementary Figure S4). Other highly expressed marker genes for C3 included cell adhesion and migration genes such as *LUM* (Lumican)*, TGFBI* (transforming growth factor beta induced), and *GABRP* (gamma-aminobutyric acid type A receptor subunit Pi).Overall marker genes were slightly enriched for “mesenchyme development” in GO analysis (Figure 2D), indicating that cells in this cluster had mesodermal characteristics, and therefore, C3 was referred to as the “mesodermal-like 1 (meso-like-1)” cluster. C4 exhibited a slightly more epithelial-like profile, with low levels of epithelial stem cell genes such as *TP63* (Figure 2C; Supplementary Figure S4) that gave rise to a slight enrichment for “epidermis development” in GO analysis. Therefore, we named C4 as the “early epithelial (early-epi)” cluster. Furthermore, marker genes for both clusters were significantly enriched in “cell-substrate adhesion”, “tissue migration” and “wound healing” associated GO-terms (Figure 2D). Consistent with the scRNA-seq data, IF of the D10 cell culture samples demonstrated wide expression of ABCG2, with p63α coexpression in a subset of ABCG2-positive cells (Figure 3B).

Both C5 and C6 displayed epithelial characteristics. C5 expressed high amounts of *TP63* as well as other epithelial and known LSC genes including *KRT14* and *CXCL14* (CXC motif chemokine ligand 14) (Figure 2C; Supplementary Figure S4). C5 marker genes were highly enriched for “epidermal development” and “skin development” GO terms, indicating the epithelial stem cell feature of this cluster. Therefore, we annotated C5 as the “induced limbal stem cell 1 (iLSC-1)”. Marker genes for C6 included epithelial genes, such as *AQP3* (aquaporin 3)*, MUC16* (mucin-16) and *S100A9* (S100 calcium binding protein A9), which were enriched for GO terms such as “epidermal differentiation” (Figure 2D), indicating cells with early stratification characteristics that are mostly studied in the skin (hence the GO analysis link to epidermis). Low expression of skin (*KRT1*) and hair follicle *(KRT72)* associated keratins (Moll et al., 2008) was detected in a few cells (Supplementary Figure S4). Nevertheless, we annotated C6 as “induced limbal stem cell 2 (iLSC-2)” cluster.

IF of D24 cell cultures confirmed protein-level expression of C5/iLSC-1 markers CK14, CXCL14, p63α/p40 (ΔNp63 isoform of p63, together indicating the most cornea-associated isoform ΔNp63α, Di Iorio et al. 2005), as well as another limbal cytokeratin CK15 (encoded by *KRT15*) (Figure 3C). Also MUC16 was expressed in D24 cells; however, its staining in IF was mainly present in morphologically distinguishable, layered cell areas accompanied with low p63α staining intensity (Figure 3D), indicating that these cells possessed a distinct epithelial cell state corresponding to the C6/iLSC-2 cells.

Interestingly, almost no transcript of the corneal lineage marker paired box 6 (PAX6) was detected in the scRNA-seq data in any of the clusters (Supplementary Figure S4). However, our real-time quantitative PCR (RT-qPCR) analyses showed upregulated PAX6 throughout the differentiation process (Supplementary Figure S5A) and in IF, we could see low but increasing intensity of nuclear PAX6 staining in the majority of cells on D10/11 and D24 (Figure 3B-C), accompanied by areas of clearly PAX6+ cells especially in the hESC and hiPSC1 cultures (Supplementary Figure S5B). Together with epithelial markers such as p63 and CK14 (Figure 3B-C), these findings indicate successful commitment to CE lineage. Taken together, cell states of the time point-specific C1-2/PSC, C3/meso-like-1, C4/early-epi, C5/iLSC-1, and C6/iLSC-2 were identified by scRNA-seq and confirmed on the protein level in IF. Cells in these clusters seemed to follow the expected progression of differentiating hPSCs towards the corneal LSC-like cells.

### 2.4. Mixed time point populations contain non-epithelial cells which deviate from the LSC differentiation pathway

Among the mixed time point clusters, C7 and C8 cells expressed genes that are normally highly expressed in mesenchymal cells or in mesodermal cells during development (Figure 2C; Supplementary Figure S4). For example, *COL1A1* (collagen 1A1)*, ZEB2 (*zinc finger E-box binding homeobox 2*), ACTC1* (actin alpha cardiac muscle 1) and *ACTA2* (alpha-smooth muscle actin) were high in both clusters. Additionally, C7 cells expressed high levels of *COL8A1* (collagen 8A1)*, GREM1* (gremlin 1), and *TGFB2* (transforming growth factor beta 2) and C8 marker genes included *HAND1* (heart and neural crest derivatives expressed 1), *GATA4* (GATA binding protein 4), *PITX1* (paired like homeodomain 1), *TGFBI,* and *LUM*. Therefore, we annotated C7 as “mesodermal-like-2 (meso-like-2)” and C8 as “mesodermal-like-3 (meso-like-3)” cluster (Figure 2). Interestingly, some of these markers, e.g., *ACTC1, ACTA2, HAND1* and *LUM* were expressed in both C3/meso-like-1 and C8/meso-like-3, indicating a somewhat similar cell states (Figure 2C; Supplementary Figure S4). IF confirmed the expression of HAND1, GATA4, lumican and ZEB2 on the protein level, with varying co-staining patterns at D10 and D24 (Figure 4A). However, clear expression of collagen 8A1 was only observed at D24 (Figure 4A), indicating that it marks a distinct cell cluster.

**Figure 4.**
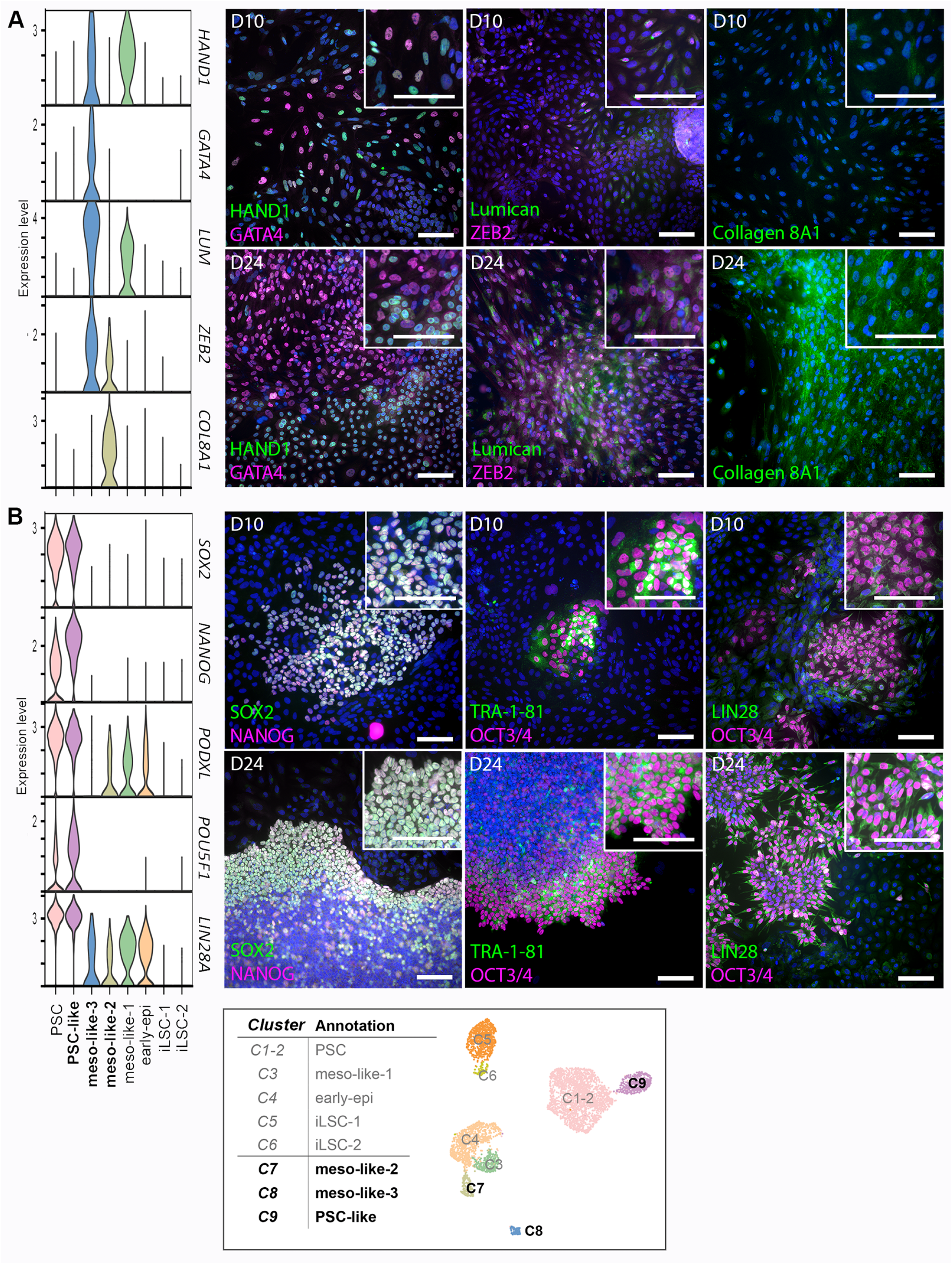
Immunofluorescence validation of mixed time point Clusters 7-9. A) Expression of Cluster 7 and/or 8 (meso-like-2 and/or -3) markers *HAND1* (HAND1), *LUM* (Lumican) *ZEB2* (ZEB2), and *COL8A1* (Collagen 8A1) at D10 and D24 time points. B) Expression of pluripotency/Cluster 9 (PSC-like) markers *SOX2* (SOX2), *NANOG* (NANOG), *PODXL* (TRA-1-81), *POU5F1* (OCT3/4), and *LIN28A* (LIN28) amongst the differentiating cells at D10 and D24 time points. Representative IF images are shown for line hiPSC1. Cell nuclei counterstained with DAPI or Hoechst 333420 (both shown in blue). Scale bars, 100 µm in all images.

Intriguingly, C9 had a high resemblance to C1-2/PSC, which represented typical undifferentiated hPSCs at D0. Similar pro-proliferation and pluripotency-associated genes were detected among the highly expressed markers in both clusters. However, one clear difference was that C9 cells expressed distinct metallothionein (*MT*) genes, which were not expressed by C1-2 (Figure 2C; Figure 4C). Nevertheless, GO-term enrichment of C9 was similar to that of C1-2 (Figure 2D), and thus this cluster was annotated as the “PSC-like” cluster. As the PSC-marker expressing cells belonging to C1-2 were only obtained at D0 (Figure 2B), we used a panel of pluripotency markers to confirm the presence of C9 cells at D10 and D24. Indeed, strong expression of SOX2, TRA-1-81, OCT3/4, and LIN28 was confirmed by IF in colonies showing undifferentiated morphology and becoming more abundant within cells at D24 when compared to D10 (Figure 4A). It was notable that, although almost no hESC-derived cells contributed to C9 in the scRNA-seq data, some formation of PSC-like colonies and expression of pluripotency markers was observed also in the D24 hESC cultures during our IF validation (Supplementary Figure S5C), although in lesser extent compared to the hiPSC1 line (Figure 4B).

### 2.5 Surface marker quantification confirms differences in the cell line-dependent differentiation efficiency

Based on the marker genes, the time point-specific cluster C5/iLSC-1 appeared to contain the most LSC-like cells. Dissection of the scRNA-seq data annotation showed that a clear majority (95%) of hESC-derived cells collected at D24 belonged to this cluster, while only 35% of hiPSC1- and 22% of hiPSC2-derived cells clustered as C5/iLSC-1 at D24 (Figure 5A). Because of the inconsistency of the two hiPSC lines, we sought to identify surface marker genes for potential purification of C5/iLSC-1. For this purpose, *AREG* (amphiregulin) and *ITGA6* (integrin subunit alpha 6) were found (Figure 5B). Furthermore, as the C9 cells constituted a prevalent but particularly undesirable population among the hiPSC lines, we aimed to specifically exclude these cells by identifying *PODXL* (encoding TRA-1-81) as the negative selection marker for LSC-like cells (Figure 5C).

**Figure 5.**
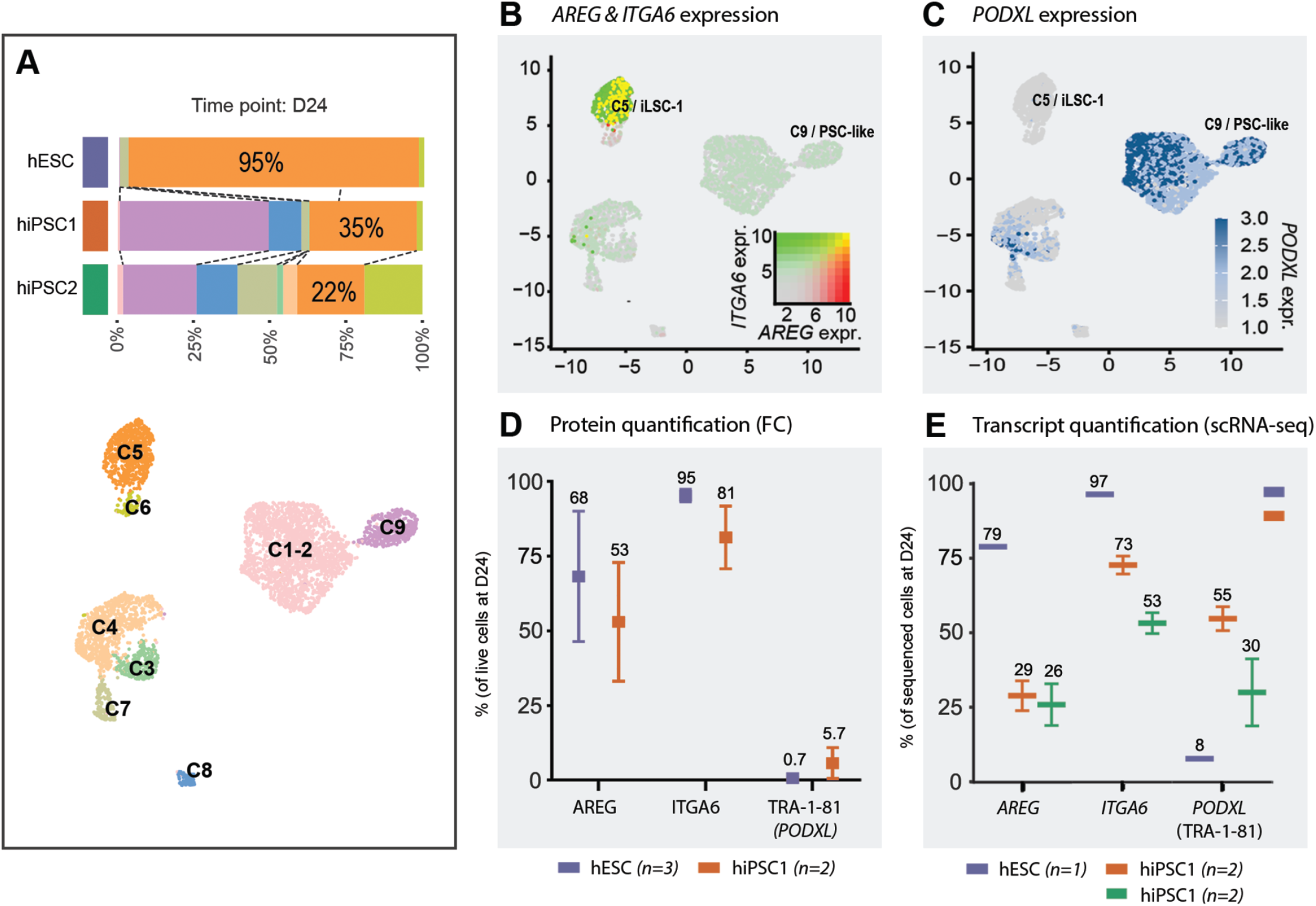
Measuring differentiation efficiency via quantification of surface markers ITGA6, AREG, and TRA-1-81 with flow cytometry (FC). A) Distribution of cells to the identified clusters at day (D)24, shown separately for each cell line. B) Expression *AREG* and *ITGA6*, and C) *PODXL* transcripts illustrated on the UMAP graphs. D) FC quantification of AREG, ITGA6, and TRA-1-81 antigens at D24. E) Quantification of cells with *AREG, ITGA6*, and *PODXL* transcripts measured in the single-cell RNA-sequencing data. Results shown as mean ± SD.

In flow cytometry (FC) analysis of hPSC-LSC populations at D24, AREG was expressed in 68.2 ± 17.8% of hESC-vs. 53.1 ± 14.1% of hiPSC1-derived cells, and ITGA6 was expressed in 95.3 ± 1.7% of hESC-vs. 81.4 ± 7.5% of hiPSC1-derived cells (Figure 5D). At the same time, TRA-1-81 was expressed only in 0.7 ± 0.4% of hESC-but in 5.7 ± 3.7% of hiPSC1-derived cells (Figure 5D). The lower amount of AREG and ITGA6 protein and higher amount of TRA-1-81 in the hiPSC1-derived cells showed the similar trend as the scRNA-seq measurements of *AREG, ITGA6,* and *PODXL* transcripts (Figure 5E): only 29% of hiPSC1 cells expressed *AREG* and 73% expressed *ITGA6* at D24, in comparison to 79% of hESC cells expressing *AREG* and 97% expressing *ITGA6.* On the other hand, *PODXL* was expressed in 55% of hiPSC1 cells and only in 8% of hESCs. To conclude, quantification of C5/iLSC-1 cells with AREG, ITGA6 and TRA-1-81 as markers confirmed more efficient derivation of C5/iLSC-1 from hESC, as compared to hiPSC1 (Figure 5D).

### 2.6 FACS-mediated selection allows a more robust generation of LSC-like cells

To test if the line-dependent differentiation efficiency can be overcome by purification of specific cell state, an *in silico* purification was simulated by computationally selecting the most LSC-like population (C5/iLSC-1) and comparing their transcriptome to data from all D24 cells and primary adult human LSCs. For this comparison, scRNA-seq data of C5/iLSC-1 cells, all D24 cells and primary adult human LSCs (Smits et al. (2023) (donor sample IDs: GSM6266908 and GSM6266909) were processed into pseudobulk data where the single cell data was aggregated, and visualized with the principle component analysis (PCA). Since 95% of the D24 hESC cells were annotated as C5/iLSC-1, but the two hiPSC lines were heterogeneous, we expected that the hESC “all D24 cells” (purple triangle) would be more similar to the “primary LSCs” (blue squares) than “all D24 cells” from hiPSC1 or hiPSC2 (orange and green triangles, respectively). We also expected that the *in silico* purification would notably improve the quality of the hiPSC lines in terms of this similarity. Indeed, this was the case, as demonstrated by positioning of the purified populations (purple/orange/green circles) closer to “primary LSCs” on the PC1 axis that represents the major difference in this PCA plot (58%) (Figure 6A).

**Figure 6.**
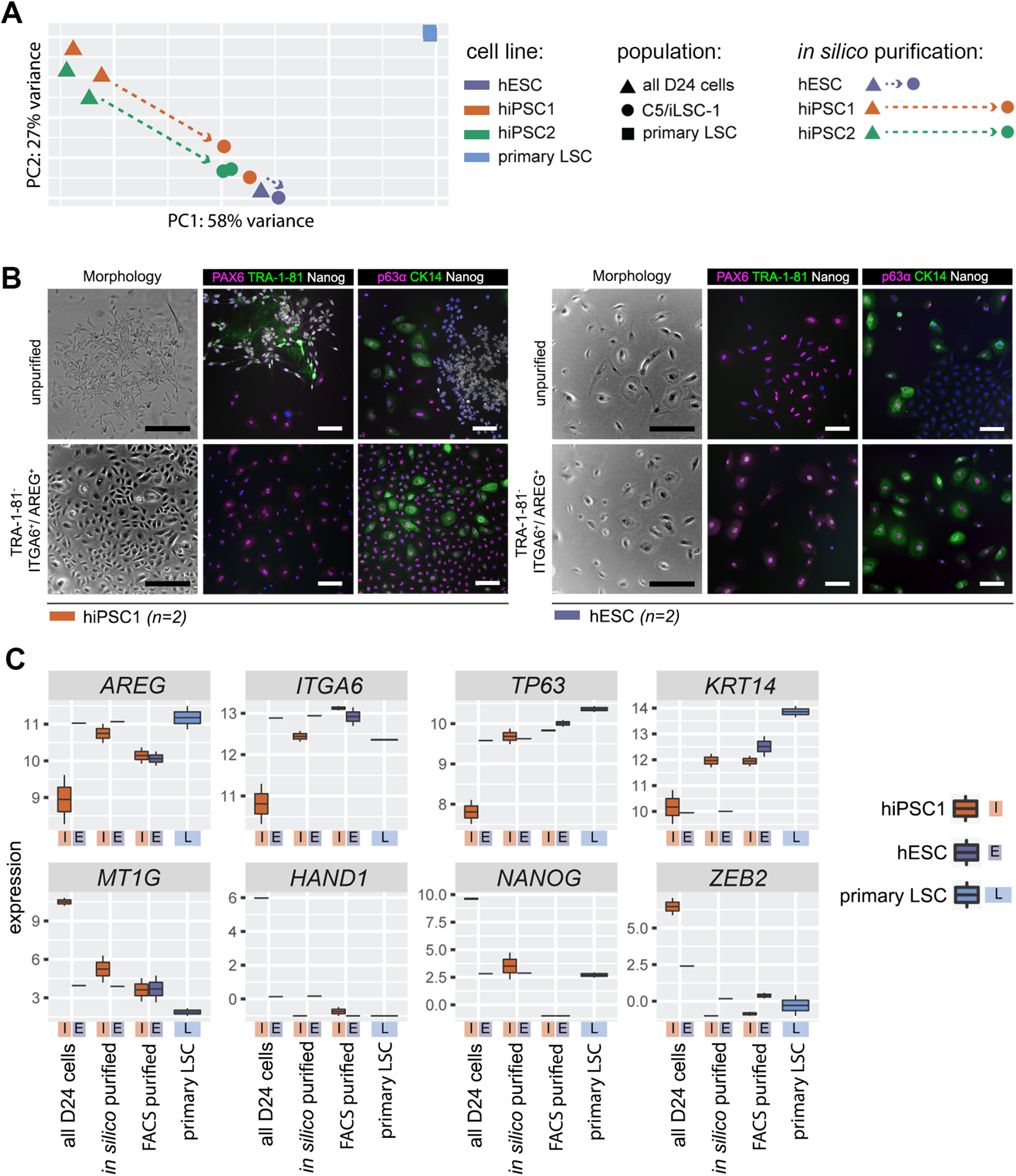
FACS-mediated purification of the target population. A) Principal component analysis (PCA) of *in silico* purified cells from the scRNA-seq data (all D24 cells vs. Cluster 5 cells) and for comparison, control data from primary cultured limbal stem cells (LSCs). B) IF characterization of unpurified vs. TRA-1-81-/ITGA6+/AREG+-purified cells at day D8 post-sorting. Cell nuclei counterstained with Hoechst 333420 (shown in blue). Scale bars: 250 µm (black) and 100 µm (white). C) Selected marker gene expression between *in silico* and FACS-purified hiPSC-derived (i) and hESC (E) populations, compared to primary LSCs (L).

We then proceeded to utilize the surface markers identified in Chapter 2.5 for actual purification of C5/iLSC-1 cells from all D24 cells through fluorescence-activated cell sorting (FACS). TRA-1-81 was to exclude C9/PSC-like cells, and ITGA6 and AREG to double-positively select Custer 5/iLSC-1 (Supplementary Figure S6A). The TRA-1-81^-^/ITGA6^+^/AREG^+^ cells and unpurified cells without any marker-based selection as the control were sorted with FACS, subcultured for 8 days, and characterized in IF (schematic overview of the workflow presented in Supplementary Figure S6B). In the post-sort cultures, multiple PSC-like colonies could be observed within the hiPSC1-derived cells without purification, whereas TRA-1-81^-^/ITGA6^+^/AREG^+^ cultures did not display any PSC-like morphology (Figure 6B). The findings were confirmed in IF, which demonstrated clear expression of NANOG and TRA-1-81 within the unpurified cells, whereas the TRA-1-81^-^/ITGA6^+^/AREG^+^ cultures expressed PAX6, p63α and CK14, but not NANOG or TRA-1-81 (Figure 6B). No PSC-like colony morphology or NANOG/TRA-1-81 expression was detected within the hESC-derived cells either with or without purification (Figure 6B). Additional IF characterization included MUC16, lumican, and CE-specific cytokeratin 12 (CK12), to reveal the potential presence of C6 (iLSC-2), C8 (meso-like-3), or terminally differentiated corneal epithelial cells, respectively, but these markers were not detected in any of the samples.

To further assess the molecular profile of the FACS-purified cells, we collected samples from TRA-1-81^-^/ITGA6^+^/AREG^+^ hESC and hiPSC1 at 15 days post-sorting and carried out bulk RNA-seq. The expression of selected cluster marker genes in the bulk RNA-seq data of FACS-purified cells were compared to the pseudobulk datasets from our “all D24 cells” and the primary LSCs from Smits et al. 2023 (Figure 6C). Similar to the *in silico* purification and supporting the IF staining results, expression of genes in FACS-purified TRA-1-81^-^/ITGA6^+^/AREG^+^ hiPSC1 cells showed more similarity to hESC “all D24 cells” and primary LSCs, as compared to all unpurified hiPSC1 “all D24 cells”, including increased expression of LSC markers and decreased expression of mesodermal-like and pluripotent markers. This indicates that our cell sorting strategy, based on surface markers identified from scRNA-seq data, can increase the hiPSC-LSC differentiation efficiency and generation C5/iLSC-1 yield, thus circumventing the line-dependent variability within the hPSCs.

## Discussion

Understanding the diverse cell types involved in the *in vitro* differentiation of hPSC is crucial to gain control over the process and advance safe and effective hPSC-based therapeutics. To achieve this, streamlined data-driven tools are ultimately needed to facilitate the development of standardization strategies and to support the production of uniform high-quality hPSC-derived cells. In this study, we showed successful differentiation of three hPSCs lines toward the LSC lineage, identifying novel cell types/states present along the process using scRNA-seq, and validating these results by IF and FC. Moreover, we demonstrated proof-of-principle that undesired populations such as non-epithelial or PSC-like cells among the differentiated LSC-like cells and high variation in the cell line-dependent differentiation efficiency can be overcome by using ITGA6 and AREG for purification of LSC-like cells. The results support our previous works (Hongisto et al., 2017, 2018; Vattulainen et al., 2019, 2021) by replicating several main findings, and more importantly, provide new insights into the differentiation heterogeneity and strategies for ensuring consistent, cell line-independent differentiation outcome.

In this study, we successfully characterized the different cell states with specific marker gene expression at different time points during the 24-day differentiation. These populations were represented by the time point-specific C1-2/PSC (marked by pluripotency genes e.g., *POU5F1* and *NANOG*), C4/early-epi (marked with early LSC genes *ABCG2* and *TP63*) and C5/iLSC-1 (marked by LSC/basal limbal genes e.g., *TP63* and *KRT14*). At the same time, we were able to identify previously unreported sub-populations containing distinct cell states. For example, C3/meso-like-1 comprising *ABCG2^+^/TP63*^-^ cells collected at D10/11 demonstrated elevated expression of genes associated with migration, invasiveness, and extracellular matrix production; indications towards epithelial-to-mesenchymal transition (EMT) (Kalluri & Weinberg, 2009). Accordingly, the upregulated markers for this cluster included mesenchymal genes like *GABRP*, *TGFBI* and *LUM*, but also a low level of early epithelial genes as well as neural crest-associated *HAND1*. While EMT does not play a major role in the surface ectoderm which develops into corneal epithelium, it is a crucial process in the closely related neural ectoderm (Weigele & Bohnsack, 2020): the interaction between the surface ectoderm and the neural ectoderm plays an important role in driving neural crest EMT and ocular morphogenesis. Interestingly, there are also indications that neural crest border cells retain plasticity to contribute to multiple ectodermal lineages (Kobayashi et al., 2020; Roellig et al., 2017). Therefore, this cluster of cells could recapitulate unintended EMT events potentially linked to developmental neural crest driven EMT. It is an interesting question warranting further investigation, whether this less-epithelial cell population represents a mere side product of differentiation, a transitional state, or a necessary factor influencing the proper differentiation of the LSC-like cells.

Although many genes in C4/early-epi were also associated with EMT, almost half of the cells also expressed epithelial *TP63*, traditional hallmark of clinically relevant LSCs (Rama et al., 2010). Moreover, while ABCG2 also serves as a stem cell marker, expression of ABCG2 together with p63 has been specifically associated with slow-cycling LSCs with high clonogenic capacity (Budak et al., 2005; de Paiva et al., 2005). We have previously demonstrated that ABCG2^+^/p63α^+^ hPSC-LSCs have enhanced regenerative abilities compared to ABCG2^-^/p63α^+^ cells both *in vitro* and *ex vivo* (Vattulainen et al., 2019). In this study, we demonstrated that D10/11 hPSC-LSCs co-express p63α and PAX6 proteins, which identify the corneal epithelial stem cells/progenitors both in tissue- and hiPSC-derived LSC cultures (Hayashi et al., 2017; Norrick et al., 2021). Therefore, we expect C4 to contain early LSC-like cells, and the precursors of the later emerging epithelial population represented by C5 (iLSC-1).

C5 expressed several traditional limbal progenitor/LSC marker genes e.g., *KRT14*, *KRT15,* and *TP63* (Di Iorio et al., 2005; Figueira et al., 2007; Nieto-Miguel et al., 2011; Pellegrini et al., 2001). It also had a high expression of a more novel marker CXCL14, which was recently associated with potential LSCs in human corneas using scRNA-seq (Català et al., 2021; Collin et al., 2021). Thus, C5 seemed to represent the most LSC-like population obtained during the differentiation process. However, the distinct profile from primary LSCs, coupled with the lack of mature corneal epithelium (CE) marker CK12 expression even after continued post-sort culture period strongly indicate that C5/iLSC-1 cells remain rather immature. Manipulation of their serum- and feeder-cell free culture environment may be required to facilitate further lineage-commitment and proper differentiation. It’s worth mentioning that in our data, PAX6 expression remained almost undetectable by scRNA-seq. However, additional analyses with RT-qPCR and IF contrasted this result by demonstrating PAX6 expression both on mRNA and protein level. This may require further attention in our upcoming studies, since correct dosage of PAX6 is known to be essential for corneal epithelial health, starting early on from the eye field development all the way to corneal morphogenesis and postnatal CE (Shaham et al., 2012; Sunny et al., 2022). Importantly, it is one of the key transcription factors driving the lineage-commitment toward the ocular epithelia instead of skin epidermis (Kitazawa et al., 2017; G. Li et al., 2015; Ouyang et al., 2014; Smits et al., 2023).

C6/iLSC-2 was identified as the slightly more differentiated epithelial population appearing at D24, characterized by e.g., MUC16, which is expressed superficially throughout the adult ocular surface epithelia (Argüeso et al., 2003) and in the developing CE (Collin et al., 2021). In our experiments, the IF staining of MUC16 was shown to localize to the apical cells of layered islands, similar to areas of immortalized corneal-limbal epithelial cell cultures (Argüeso et al., 2003). However, as the CE-specific differentiation marker *KRT12* (CK12) was not detected, cells in this cluster may also have other epithelial fates. In line with this reasoning, low expression of *KRT1* and *KRT72* keratins which were detected in a small subpopulation of iLSC-2 cells) indicated partial off-target differentiation toward epidermis or hair follicles (Moll et al., 2008).

In addition to the time point-specific C1-5, we also detected three clusters containing cells from multiple time points (C7-9). Two of these clusters, C7 and C8 contained cells from both D10/11 and D24 and displayed even higher EMT genes expression than the time point-specific C3 cells. Activation of unintended neural crest EMT gene expression programs could also explain the origin of these cells. However, unlike the cells in C3 that either go extinct during the process or develop further into a new phenotype, the cells in Clusters 7 and 8 remain and overcommit to a mesodermal gene signature. Markers expressed in these clusters included genes associated with mesodermal differentiation and often linked to cardiac development, e.g., *ACTC1, HAND1* and *GATA4* (Olson, 2006); this may indicate a diversion from the intended LSC fate.

Finally, C9 represented a population deviating even further from the intended cell type. It contained PSC-like cells from all three time points, with a large percentage of D24 cells. PSC marker expression coupled with high proliferation rate are known risks for tumorigenicity (Lee et al., 2013; Yamanaka, 2020). The distinctive attribute of C9 compared to C1-2 (PSCs) was the clear upregulation of MT genes, which play multiple roles in carcinogenesis, including regulation of cell cycle arrest, proliferation, and apoptosis (Si & Lang, 2018). Some cells in C9 were derived from D0 time point, thus showing high initial MT levels. In a study of Lu et al. (2018), expression of MTs was shown to characterize a bizarre differentiation-arrested population, which was potentially linked to altered cellular responses to medium zinc in a rare population of hESCs. It would be of interest to further investigate the mechanisms of these genes maintaining PSC-like phenotype during differentiation, and to unravel if certain hPSC line characteristics make it more prone to produce such unfavorable outcome.

In this study, the general differentiation efficiency was highest in the hESC line, with 95% of the cells belonging to the most LSC-like C5, while both hiPSC1 and hiPSC2 produced noticeably lower number of cell following the differentiation trajectory and contributed more to mixed time point Clusters 7, 8, and 9. This may not be fully surprising, given that the used protocol has been initially optimized with the hESC line (Hongisto et al., 2017). The outcome is most likely also influenced by the differences in cell line origin: embryonic or reprogrammed, and even by the reprogramming method (Bock et al., 2011; Cahan & Daley, 2013; Osafune et al., 2008; Yamanaka, 2020). However, the observed differences represent the well-acknowledged problems of line-dependent hPSC differentiation efficiency. This may also explain the challenges to reproduce the previously reported differentiation efficiencies with different cell lines, despite using the same protocol (Sun et al., 2021).

To solve these issues, scRNA-seq data-driven decisions can be made to further optimize the differentiation protocol for each individual cell line. However, this approach is time-consuming and laborious, when considering its economic feasibility for translational purposes. As implicated by the pseudobulk RNAseq data of the *in silico* purified C5 cells, selective enrichment could be used to efficiently mitigate the effect of lower differentiation efficiency and inconsistency between different hPSC lines. In this study, we successfully demonstrate this proof-of-principle: by combining positive selection of C5-specific surface markers AREG and ITGA6 with negative selection of pluripotency marker TRA-1-81 in FACS-mediated selection the target population, we efficiently purified C5 cells and removed all other cells, including the potentially high-risk PSC-like cells. Ultimately, the selective sorting evened out the difference between the golden standard hESC line and the less efficient hiPSC1, by significantly enhancing the post-sort efficiency of hiPSC1 in terms of LSC-associated gene and protein level expression. When compared to the gene expression of primary adult LSCs, the purification was shown to further improve the quality of the hPSC-derived cell populations based on the elevated expression of important LSC marker genes such as *TP63* and *KRT14*, and decreased level of off-target markers like *HAND1* and *ZEB2*. To the best of our knowledge, our study represents the first to employ scRNA-seq for a detailed investigation of differentiating hPSC-LSCs heterogeneity, to define the precise cell states, differentiation efficiency and cell line-dependency. Based on specific markers identified by scRNA-seq, we successfully improve the quality and consistency of the obtained hPSC-LSCs through cell sorting. Importantly, such data-driven strategy can be widely applicable to any hPSC-based cell production, increasing the chances of using these cells in clinics.

## 3. Experimental procedures

### 3.1. Corneal differentiation of hPSCs

Three genetically distinct hPSC lines were used in this study. Human ESC line Regea08/017 (“hESC”) was derived and characterized as described by Skottman (2010). Human iPSC line WT001.TAU.bB2 (“hiPSC1”) was established and characterized in-house as described in Grönroos et al. (2021), and the other hiPSC line WT003.TAU.bC (“hiPSC2”) as described in Supplementary File 1. Corneal differentiation of hPSCs for 24 days was performed with established protocol as previously described in full detail by Hongisto et al. (2017, 2018) and Vattulainen et al. (2019, 2021).

### 3.2. Single-cell RNA-sequencing (scRNA-seq)

Cell cultures at D0, D10/11, and D24 were enzymatically dissociated to single cell suspension and stained with BD Horizon™ Fixable Viability Stain 510 (FVS510, BD Biosciences). Live single cells were sorted onto 384-wells containing primers with unique molecular identifiers (UMIs), according to the SORT-Seq protocol (Hashimshony et al., 2016; Muraro et al., 2016). Sorting was performed with BD FACSAria™ Fusion cell sorter, operating with BD FACSDiva™ v8.0.1 software (both from BD Biosciences). D0, D10/11 and D24 samples were collected from two individual differentiation batches of both hiPSC lines and one differentiation batch of the hESC.

For the processing of the samples and construction of the libraries, External RNA Control Consortium (ERCC) spike-in mix (1:50 000) was added to the wells by Nanodrop (BioNex Inc), followed by the addition of 150 nl Reverse transcription mix. After thermal cycling (4 °C 5 min; 25 °C 10 min; 42 °C 1 h; 70 °C 10 min), the plate libraries were pooled together and AmpureXP beads (New England BioLabs) were used to purify the cDNA. *In vitro* transcription with MEGAscript™ (Invitrogen) was carried out overnight at 16 °C, with the lid set at 70 °C.

An exonuclease digestion step was thereafter performed for 20 min at 37 °C, followed by fragmentation of the RNA. Another AmpureXP bead cleanup was performed to purify the RNA, after which the samples were subjected to library reverse transcriptase and amplification to tag the RNA molecules with specific and unique sample indexes (Illumina), followed by a final beads cleanup (1:0.8, reaction mix: beads) and elution of sample cDNA libraries with DNAse free water. Libraries were quantified using the KAPPA quantification kit following manufacturers protocol, after which the plates were sequenced on the NextSeq 500 (Illumina) for 25 million reads per plate.

### 3.3. Data processing

Single cell libraries were pre-processed using seqscience (Van Der Sande et al., 2023). Briefly, reads were aligned using kalistobuss to GRCh38.p13. Cells underwent QC filtration using Seurat, which involved excluding cells with low UMI counts, low measured genes, high ERCC spike-ins, and/or contaminations (with DecontX; Yang et al., 2020) as well as potential doublets identified using doubletFinder (McGinnis et al., 2019). (Supplementary Figure S2A). Data variance was assessed using Scater (Supplementary Figure S2B-C) which indicates the time point of cells drove most of the variance within the data. Seurat 4 was used for analysis (Hao et al., 2021). PCs were selected for dimensionality reduction (Supplementary Figure S2D-E), after which UMAP dimensionality reduction was performed followed by Louvain clustering (Supplementary Figure S3A). Cluster resolution stability was assessed using clusttree (Zappia & Oshlack, 2018). This resulted in an extensive robust sampling of hundreds of cells across each time point, cell line, and replica (Supplementary Figure S2A; Supplementary Table S1). Estimation of cell cycle state was performed using Seurat CellCycleScoring() feature with the cell cycle genes from Tirosh et al. (2016) (Figure 2B-C, Supplementary Figure S3). Finally, the QC values were compared between clusters (Supplementary Figure S3E).

### 3.4 Marker gene selection and GO-term enrichment

Marker genes for each cluster were identified using the Wilcoxon Rank Sum test (Stuart et al., 2019). GO-term enrichment was performed using separated positive and negative marker genes for each cluster with clusterprofiler (Wu et al., 2021). Finally, a combination of marker genes, GO-term enrichment, and sample time point information was used to generate cluster annotations. The comprehensive gene lists for each cluster are also provided in Supplementary File 4.

### 3.5 Real-time quantitative PCR (RT-qPCR)

The RT-qPCR analysis was performed on three individual differentiation batches of hESC and two batches of both hiPSC1 and hiPSC2. RNA was isolated from cell pellet samples with Qiagen RNeasy® Plus Kit (ThermoFisher). After removal of genomic DNA and synthetization of complementary DNA, all samples were run as triplicate reactions with the Applied Biosystems QuantStudio 12K Flex Real-Time PCR system using sequence-specific TaqMan™ Gene Expression Assays (ThermoFisher) for GAPDH (Hs99999905_m1) and PAX6 (Hs01088112_m1) and analyzed with −2ΔΔCt method (Livak & Schmittgen, 2001).

### 3.6 Immunofluorescence (IF)

The markers for IF validation were chosen based on their average log fold-change (avg_log2FC) expression and abundance in the target cluster vs. all the other clusters, and antibody availability. Indirect IF protocol was performed on cell cultures fixed with 4% paraformaldehyde (Electron Microscopy Sciences) as previously described (Vattulainen et al., 2021), using primary antibodies listed in Supplementary Table S2. Olympus IX51 fluorescence microscope (Olympus, Corporation) was used to capture the staining results, and representative IF image panels were constructed and edited in Adobe Photoshop 2023 (version 27.4.2, Adobe Inc.).

### 3.7 Flow cytometry FC) and sorting (FACS)

Suitable markers were screened from C5 marker genes linked to the GO-term “cell surface” (GO:0009986) based on their avg_log2FC expression and abundance in the C5 vs. all the other clusters. For FC, single cell suspension was first stained with FVS510, and the nonspecific antibody binding sites were blocked with 2,5-3% bovine serum albumin for 10-15 min at room temperature. Samples were stained for 30 min on ice with pre-optimized concentrations of the following fluorochrome-conjugated antibodies: AREG-APC (17-5370-42; Invitrogen), CD49f (ITGA6)-PE (561894/555736) and TRA-1-81 (560194; both BD Biosciences). Samples were analyzed with BD FACSAria™ Fusion, using an appropriate panel of single-stained and fluorescence minus one (FMO) controls for compensation and correct gating of the data (Maecker & Trotter, 2006) (Supplementary Figure S6A). 10 000 single cell events per sample were recorded for analysis, followed by sorting of TRA-1-81^-^/ITGA6^+^/AREG^+^ cells onto Collagen IV/laminin 521 coated wells for subculturing in Cnt-30. Live cells with no selection were sorted as controls, at a similar seeding density (10 000 - 40 000 cells per 24-well, area=1.9 cm^2^). 10 µM Rho-kinase inhibitor Y-27632 and 2% KnockOut™ SR XenoFree Medium Supplement (Gibco) were added to the medium for the first day. The optimized analysis was first tested with D24 hESC-LSCs without sorting, followed by two replicate sorting experiments with both hESC- and hiPSC1-LSCs.

## Supporting information

Supplementary File 1 003b characterization

Supplementary File 4 Marker genes

## Data availability

All raw sequencing files generated in this study have been deposited in the Gene Expression Omnibus database with the accession number GSE248497. All code used in this study is available at https://github.com/JGASmits/Heterogeneity-of-differentiating-hPSC-derived-corneal-limbal-stem-cells-through-scRNAseq/. Public data of scRNA-seq data from adult LSCs was downloaded from GSM6266908 & GSM6266909. The marker gene lists generated for identified clusters are shared in Supplementary File 4 within this manuscript.

## Abbreviations

ABCG2: ATP-binding cassette subfamily G member 2
ACTA2: alpha-smooth muscle actin
ACTC1: actin alpha, cardiac muscle 1
AQP3: aquaporin 3
CE: corneal epithelium
CK: cytokeratin
COL: collagen
CXCL14: CXC motif chemokine ligand 14
DAPI: 4′,6-diamidino-2-phenylindole
ERCC: external RNA control consortium
FACS: fluorescence-activated cell sorting
FC: flow cytometry
FMO: fluorescence minus one
GABRP: gamma-aminobutyric acid type A receptor subunit Pi
GO: Gene Ontology
GATA4: GATA binding protein 4
GREM1: gremlin 1
HAND1: heart and neural crest derivatives expressed 1
hESC: human embryonic stem cell
hiPSC: human induced pluripotent stem cell
(h)PSC: (human) pluripotent stem cells (both hESCs and hiPSCs)
IF: immunofluorescence
KRT: keratin
LSC: limbal stem cell
LSCD: limbal stem cell deficiency
LUM: lumican
MT: metallothionein
MUC16: Mucin-16
NANOG: Nanog homeobox
OCT3/4: octamer-binding transcription factor 3/4
p63 / TP63: tumor protein p63
PAX6: paired box 6
PC: principal component
PITX1: paired like homeodomain 1
PODXL: podocalyxin
POU5F1: POU class 5 homeobox 1
QC: quality control
RT-qPCR: real-time quantitative PCR
scRNA-seq: single-cell RNA-sequencing
S100A9: S100 calcium binding protein A9
SOX2: SRY-box transcription factor 2
TGFB2: transforming growth factor beta 2
TGFBI: transforming growth factor beta induced
UMAP: uniform manifold approximation and projection
UMI: unique molecular identifier
ZEB2: zinc finger E-box binding homeobox 2

## Author contribution

Conceptualization: H.Z. and H.S.

Funding, resources and overall project administration: H.Z. and H.S.

scRNA-seq sample collection, IF, FC/FACS, and cell culture: M.V.

scRNA-seq sample preparation, data curation and formal analysis: J.G.A.S.

Visualization: M.V. and J.G.A.S.

Writing – original draft: M.V. and J.G.A.S.

Writing – review and editing: M.V., J.G.A.S., T.I., D.L.C., H.Z. and H.S.

## Acknowledgements

The authors wish to thank biomedical laboratory technicians Outi Melin and Hanna Pekkanen for their technical assistance and contributions to cell culture and scRNA-seq sample sorting, and MSc student Sonja Harjuntausta for carrying out *PAX6* RT-qPCR and IF for the hESC1 and hESC2 lines. Tampere University Imaging Core and Flow Cytometry Core are thanked for providing technical assistance and equipment. Professor Aalto-Setälä group at Tampere University are acknowledged for the iPSC reprogramming plasmids. M.P.A. Baltissen, L.A. Lamers, and S. Rinzema are thanked for operating the Illumina analyzer and performing data demultiplexing.

The authors also wish to thank following funding sources for their financial support to this study: Research Council of Finland (to H.S.), Sigrid Jusélius Foundation (to H.S.), The National Eye and Tissue Bank Foundation (to M.V.), Mary & Georg Ehrnrooth Foundation (to M.V.), Ella & Georg Ehrnrooth Foundation (to M.V.), Evald and Hilla Nissi Foundation (to M.V.), Aard-en Levenswetenschappen, Nederlandse Organisatie voor Wetenschappelijk Onderzoek (to H.Z.), the European Joint Programme Rare diseases (to H.Z.), and ZonMw Open (to H.Z.). In addition, this study is based upon the work from COST Action CA18116, “ANIRIDIA-NET”, supported by COST (European Cooperation in Science and Technology).

## Supplementary Materials

**Supplementary File 1.docx:** Establishment and characterization of the hiPSC line WT003.TAU.bC

**Supplementary File 2.docx:** Supplementary Figures S1-S6.

**Supplementary File 3.docx:** Supplementary Tables S1-2.

**Supplementary File 4.xlsx:** Cluster marker gene lists.

