## Supplementary File 1 003b characterization for "Deciphering the heterogeneity of differentiating hPSC-derived corneal limbal stem cells through single-cell RNA-sequencing"

**Establishment and characterization of the hiPSC line WT003.TAU.bC**

The research group has been granted supportive statements from the Regional Ethics Committee of the Expert Responsibility area of Tampere University Hospital to establish and use hiPSC lines in ophthalmic research purposes (R16116). Herein, the establishment and characterization of the new hiPSC line WT003.TAU.bC is described. All the experiments were performed in accordance with relevant guidelines and regulations.

Peripheral blood monocytes (PBMC) were isolated from 20 mL of healthy donor blood sample (written consent obtained) and reprogrammed into human iPSCs using the 4D-Nucleofector™ System (Lonza, Basel, Switzerland) according to the manufacturer’s protocols and employing a cocktail of following plasmids (available on Addgene repository upon request) in equimolar (1:1:1:1) amounts: 27,077 (pCXLE-hOCT3/4-shp53-F); 27,078 (pCXLE-hSK); 27,080 (pCXLE-hUL) and 37,624 (pCXWB-EBNA1). Program [FI-115] was used, with 1 µg plasmid DNA cocktail for 2x 10e6 cells recovered from cryostorage 2 h prior the transfection.

After transfection, the cells were immediately plated down on laminin 521 (L521; from Biolamina) into RPMI 1640 Medium (Thermo Fisher Scientific) supplemented with 5% human serum. During the following days, RPMI 1640 was gradually replaced with N2B27-medium consisting of Dulbecco's Modified Eagle Medium/Nutrient Mixture F-12 (with HEPES) base supplemented with 1% N-2 supplement, 1% B27 supplement, 1% non-essential amino acids, 0.1 mM 2-mercaptoethanol (all from Thermo Fisher Scientific) and 100 ng/ml basic fibroblast growth factor (from Peprotech). After approximately one week culture in N2B27-medium, the medium was again gradually replaced this time with Essential 8 Flex™ (E8 Flex; Thermo Fisher Scientific). The clone used for generation of this cell line was manually selected 17 days after the transfection.

Subculture and cryopreservation steps (from passage 3 onwards) were conducted to obtain adequate master and working cell banks for research purposes, with the removal of reprogramming vectors, normal karyotype and negativity to mycoplasma confirmed (data not illustrated). The newly established cell line was characterized for its pluripotency-associated marker expression and multi-lineage differentiation capacity. The results were analyzed with immunofluorescence, using specific primary antibodies TRA-1-81, SSEA-4, NANOG, LIN28 and OCT3/4 for pluripotency and AFP, α-SMA and OTX2 for the differentiation to endoderm, mesoderm, and ectoderm, respectively. The staining results and the details of the used antibodies are presented in the net page.

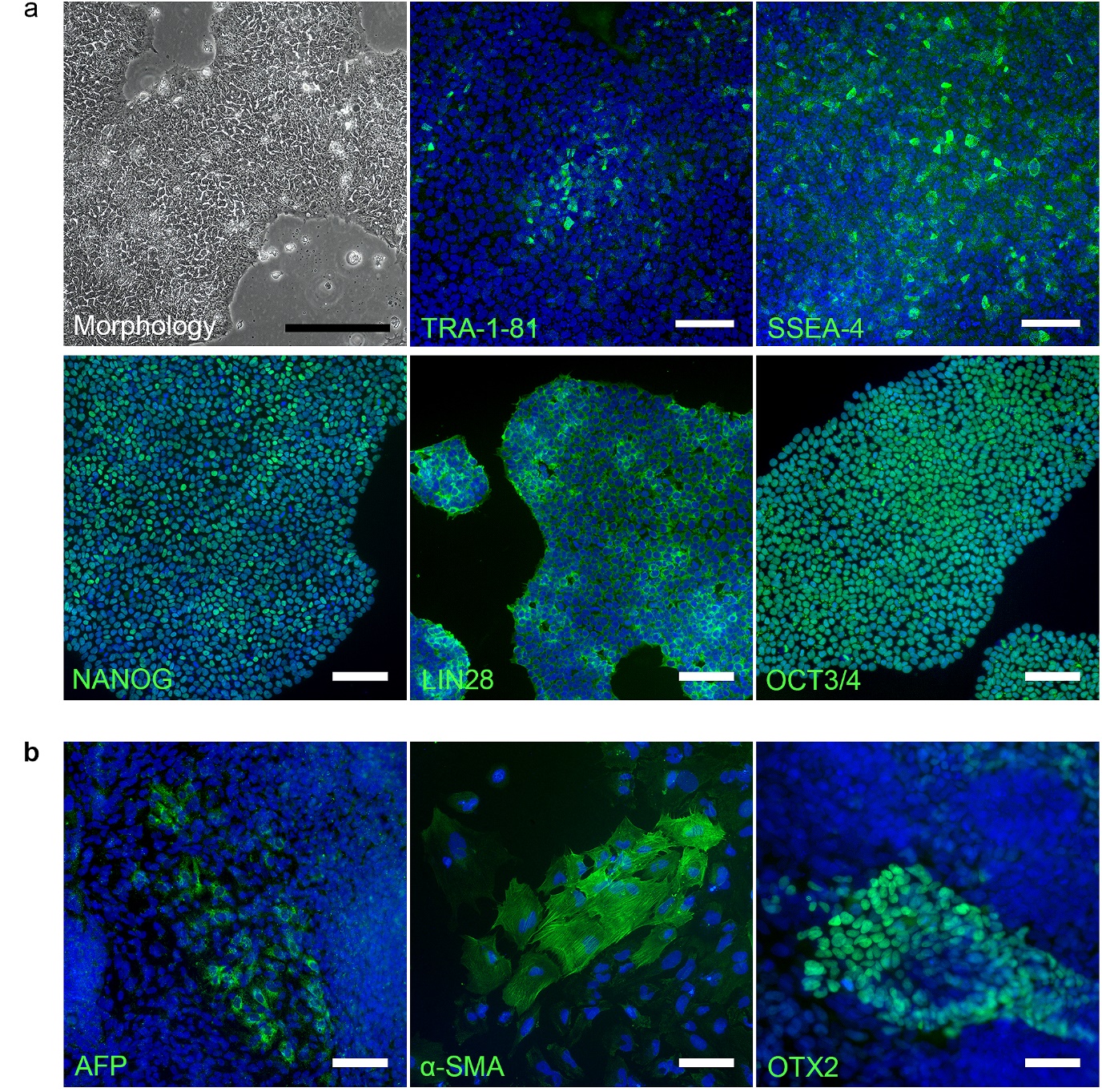

**Supplementary Results: Characterization of WT003.TAU.bC hiPSCs.** a) Colony morphology and pluripotency marker expression at passage 20. b) Differentiation towards all embryonic germ layers after spontaneous differentiation as embryoid bodies was confirmed with AFP for endoderm, α-SMA for mesoderm, and OTX2 for endoderm at passage 21. Cell nuclei counterstained with DAPI (shown in blue). Black scalebar, 200 µm. White scalebars, 100 µm.

**Supplementary Materials: Antibody information.**

|  | Antibody | Host | Dilution | Manufacturer, #Cat.No |
| --- | --- | --- | --- | --- |
| Pluripotency  markers | **TRA-1-81** | mouse | 1:400 | Santa Cruz Biotechnology, #SC-21706 |
|  | **SSEA-4** | mouse | 1:200 | R&D Systems, #MAB1435 |
|  | **NANOG** | goat | 1:100 | R&D Systems, #AF1997 |
|  | **LIN28** | mouse | 1:400 | ThermoFisher, #MA1-016 |
|  | **OCT3/4** | goat | 1:200 | R&D Systems, #AF1759 |
| Differentiation  markers | **AFP** | mouse | 1:200 | R&D Systems, #MAB1369 |
|  | **α-SMA** | mouse | 1:400 | R&D Systems, #MAB1420 |
|  | **OTX2** | goat | 1:200 | R%D Systems, #AF1979 |
