## Supplementary File 2 Supplementary Figures for "Deciphering the heterogeneity of differentiating hPSC-derived corneal limbal stem cells through single-cell RNA-sequencing"

**Supplementary Results: Supplementary Figures S1-S6**

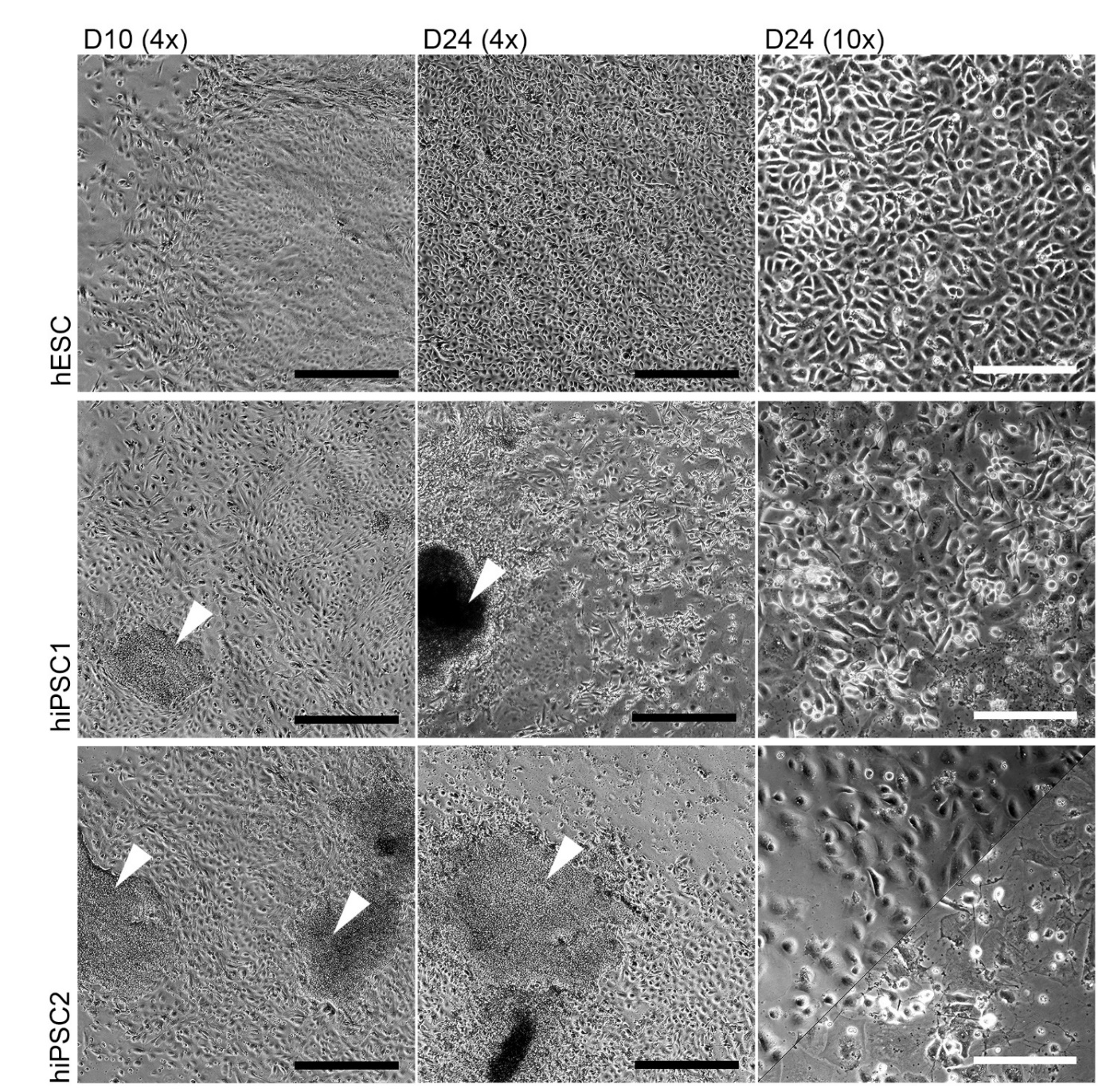

**Supplementary Figure S1. hPSC-LSC morphology during differentiation.** Representative morphologies of hESC (Regea08/017), hiPSC1 (WT001.TAU.bB2) and hiPSC2 (WT003.TAU.bC) at D10 and D24 time points during differentiation towards hPSC-LSCs. Notably lower differentiation efficiency, with more contaminating fibroblastic and/or hPSC-like colony formation was observed within hiPSC1 and hiPSC2 compared to the hESC, both during and at the end of the 24-day differentiation period. Scale bars, 500 µm (black) in 4x images and 200 µm (white) in 10x images. White arrowheads indicate hPSC-like colonies/morphologically atypical areas.

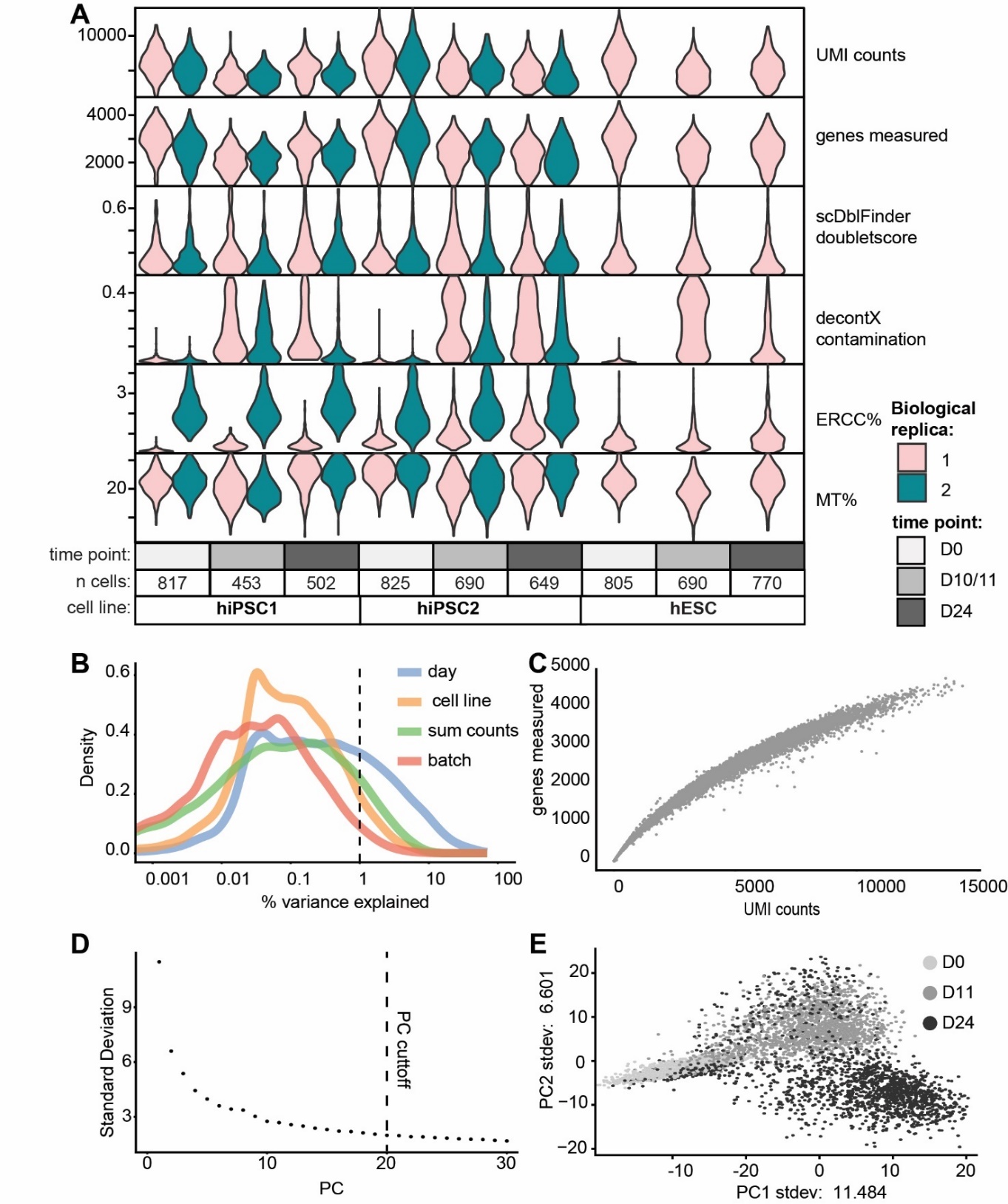

**Supplementary Figure S2. Single-cell RNA-sequencing quality control (QC) analysis.** A) QC of filtered cells. The biological replicas of each cell line are split. Violin plots of the UMI counts per cell, genes measured per cell, scDblFinder doublet score per cell, decontX contamination score, Percentage of ERCC reads, and percentage of mitochondrial reads are all plotted on the y-axis. B) Scater variance per metadata, with amount of cell density on the y-axis, and percentage of variance explained on the x axis. C) Genes measured per cell on the y-axis and UMI count measured per cell on the x-axis. D) Principal components (PC) used in dimensionality reduction. E) PC plot of PC 1 and PC2, annotated per time point.

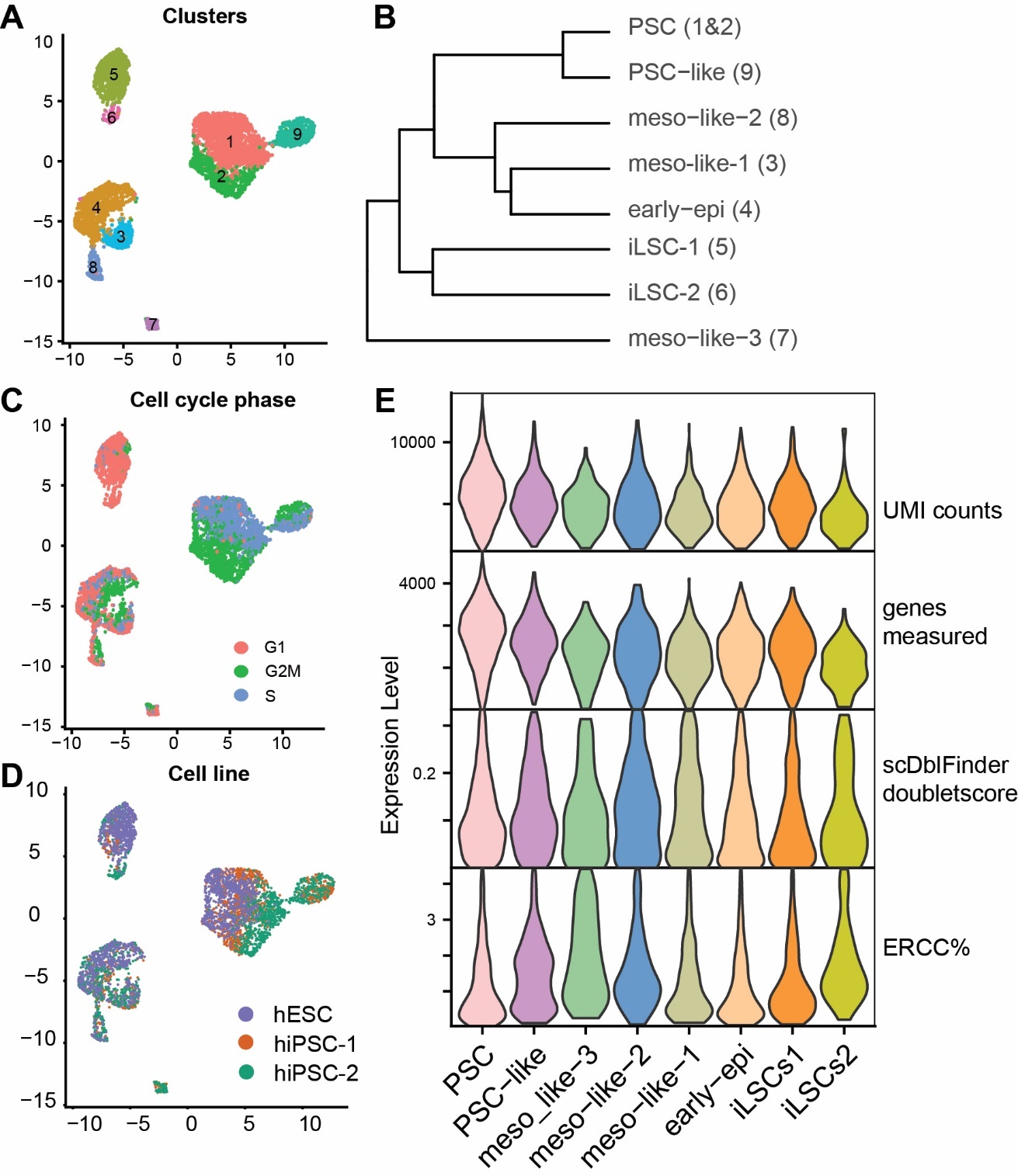

**Supplementary Figure S3. Single-cell RNA-sequencing clustering and cluster quality control (QC) analysis.** A) UMAP annotated with the clusters identified by Louvain clustering, cluster 1 and 2 were merged for subsequent analysis. B) Hierarchical tree of cluster similarity analysis C) UMAP annotated with the inffered cell cycle phase per cell D) UMAP annotated with the cell line E) Violin comparing the QC values of the annotated single cell clusters.

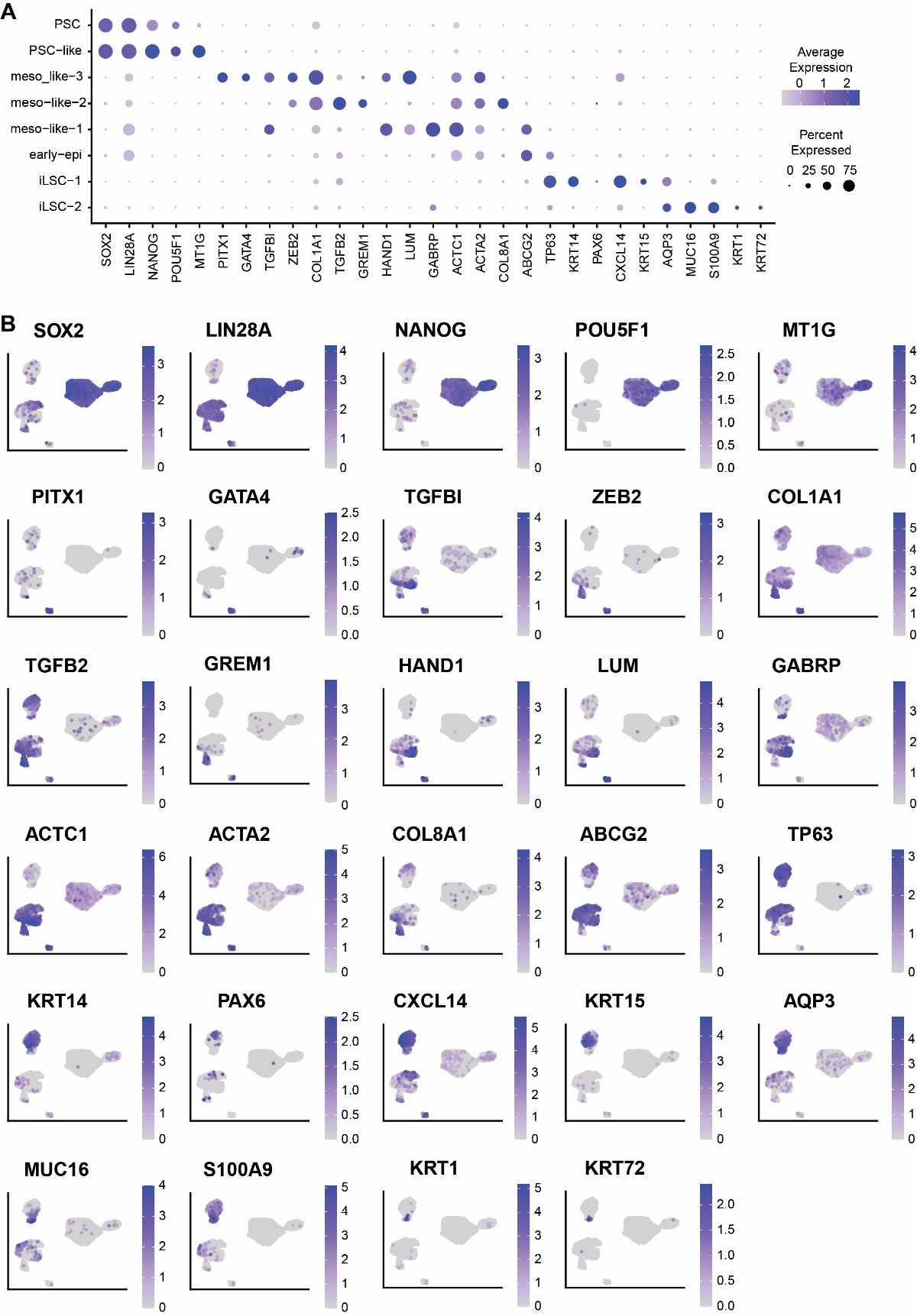

**Supplementary Figure S4. Single-cell RNA-sequencing clustering marker gene expression.** A) Expression of various cluster marker genes illustrated in the dot plot. B) The expression of marker genes plotted on top of the UMAP. Cells with the highest expression are plotted on top to better visualize low expressed marker genes.

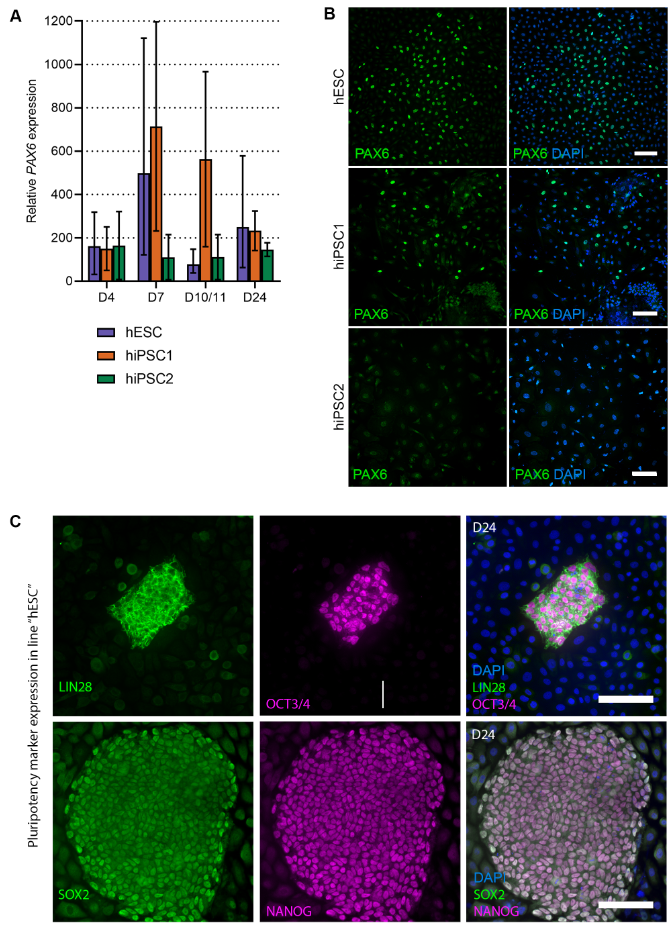

**Supplementary Figure S5**: **Expression of PAX6 and pluripotency markers during the differentiation of hPSCs towards LSCs.** A) RT-qPCR results showing the development of *PAX6* gene expression through D4, D7, D10/11, and D24 time points in hESC (Regea08/017), hiPSC1 (WT001.TAU.bB2) and hiPSC2 (WT003.TAU.bC) cell lines, presented as relative to the respective undifferentiated hPSCs at D0. B) PAX6 protein staining in hESC, hiPSC1 and hiPSC2 cell lines at D24. c) Expression of pluripotency markers LIN28, OCT3/4, SOX2, and NANOG in the PSC-like cell colonies in line hESC at D24. Scale bars for all images, 100 µm.

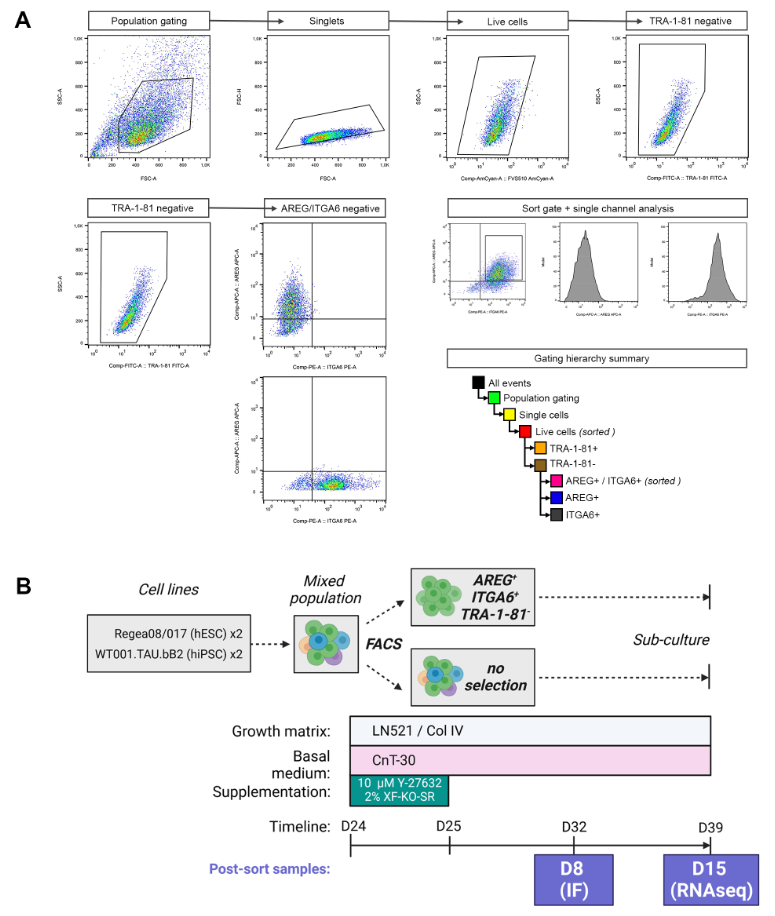

**Supplementary Figure S6. Purification of the LSC-like cells at day 24 through fluorescence activated cell sorting (FACS).** A) Gating strategy and population hierarchy summary for FACS. B) Schematic overview of the FACS experiments and the post-sort analyses.
