## Supplementary File 3 Supplementary Tables for "Deciphering the heterogeneity of differentiating hPSC-derived corneal limbal stem cells through single-cell RNA-sequencing"

**Supplementary Table S1:** Distribution of cells in clusters per time point, cell line, and cell cycle phase. Showing the number of cells and the percentage of cells from that cluster. For a visual overview, see Figure 2B.

| Cluster | Total cell number | Time point | | | Cell line | | | Cell cycle phase | | |
| --- | --- | --- | --- | --- | --- | --- | --- | --- | --- | --- |
|  |  | **D0** | **D10/11** | **D24** | **hESC** | **hiPSC1** | **hiPSC2** | **G1** | **G2M** | **S** |
| 1/2 | *n=2135* | 99.0%  *(n=2113)* | 0.6%  *(n=13)* | 0.4%  *(n=9)* | 34.6% *(n=738)* | 35%  *(n=748)* | 30.4%  *(n=649)* | 3%  *(n=64)* | 45.2%  *(n=964)* | 51.9%  *(n=1107)* |
| 3 | *n=305* | 0%  *(n=0)* | 97%  *(n=296)* | 3%  *(n=9)* | 24.3%  *(n=74)* | 34.4%  *(n=105)* | 41.3%  *(n=126)* | 35.1%  *(n=107)* | 38.7%  *(n=118)* | 26.2%  *(n=80)* |
| 4 | *n=895* | 0.1%  *(n=1)* | 97.9%  *(n=876)* | 2%  *(n=18)* | 47.9%  *(n=429)* | 18.3%  *(n=164)* | 33.7%  *(n=302)* | 49.6%  *(n=444)* | 26.4%  *(n=236)* | 24%  *(n=215)* |
| 5 | *n=735* | 0%  *(n=0)* | 0%  *(n=0)* | 100%  *(n=735)* | 73.3%  *(n=539)* | 14.8%  *(n=109)* | 11.8%  *(n=87)* | 92.5%  *(n=680)* | 3.4%  *(n=25)* | 4.1%  *(n=30)* |
| 6 | *n=91* | 0%  *(n=0)* | 0%  *(n=0)* | 100%  *(n=91)* | 11%  *(n=10)* | 6.6%  *(n=6)* | 82.4%  *(n=75)* | 97.8%  *(n=81)* | 1.1%  *(n=1)* | 1.1%  *(n=1)* |
| 7 | *n=164* | 0%  *(n=0)* | 54.9%  *(n=90)* | 45.1%  *(n=74)* | 20.1%  *(n=33)* | 15.9%  *(n=26)* | 64%  *(n=105)* | 52.4%  *(n=86)* | 34.1%  *(n=56)* | 13.4%  *(n=22)* |
| 8 | *n=105* | 0%  *(n=0)* | 18.1%  *(n=19)* | 81.9%  *(n=86)* | 0%  *(n=0)* | 38.1%  *(n=40)* | 61.9%  *(n=65)* | 71.4%  *(n=75)* | 19%  *(n=20)* | 9.5%  *(n=10)* |
| 9 | *n=410* | 21.7%  *(n=89)* | 18%  *(n=74)* | 60.2%  *(n=247)* | 0.5%  *(n=2)* | 46.8%  *(n=192)* | 52.7%  *(n=216)* | 3.9%  *(n=16)* | 38.3%  *(n=157)* | 57.8%  *(n=237)* |
| Total cell number | *n=4840* | *n=2203* | *n=1368* | *n=1269* | *n=1825* | *n=1390* | *n=1625* | *n=1561* | *n=1577* | *n=1702* |

Abbreviations: D: Day, hESC: human embryonic stem cell; hiPSC: human induced pluripotent stem cell

**Supplementary Table S2:** Immunofluorescence primary antibody information.

| **Antibody:** | **Marker gene:** | **Host:** | **Dilution:** | **Manufacturer, #Cat.No:** |
| --- | --- | --- | --- | --- |
| ABCG2 | *ABCG2* | mouse | 1:200 | Millipore, #MAB4155 |
| CK14 | *KRT14* | mouse | 1:200 | R&D Systems, #MAB3164 |
| CK15 | *KRT15* | mouse | 1:200 | ThermoFisher, #MS-1068-P1 |
| Collagen 8A1 | *COL8A1* | rabbit | 1:200 | Novus Biologicals, #25900002 |
| Collagen 8A1 | *COL8A1* | rabbit | 1:200 | Invitrogen, #PA5-97604 |
| CXCL14 | *CXCL14* | mouse | 1:200 | Invitrogen, #PA5-106402 |
| GATA4 | *GATA4* | goat | 1:200 | R&D Systems, #AF2606 |
| GATA4 | *GATA4* | mouse | 1:200 | Cell Signaling Technology, #36966S |
| HAND1 | *HAND1* | mouse | 1:50-1:100 | Santa Cruz Biotechnology, #sc-39037 |
| LIN28 | *LIN28* | mouse | 1:400 | ThermoFischer, #MA1-016 |
| Lumican | *LUM* | goat | 1:200 | R&D Systems, #AF2846 |
| MUC16 | *MUC16* | mouse | 1:100 | Abcam, #ab693 |
| Nanog | *NANOG* | goat | 1:100 | R&D Systems, #AF1997 |
| OCT3/4 | *POU5F1* | goat | 1:200 | R&D Systems, #AF1759 |
| p40/ΔNP63 | *TP63* | mouse | 1:100 | BioCare Medical, #ACI3066A |
| p63a | *TP63* | rabbit | 1:100-1:200 | Cell Signaling Technologies, #4892 |
| PAX6 | *PAX6* | rabbit | 1:200 | Sigma-Aldrich, #HPA030775 |
| SOX2 | *SOX2* | mouse | 1:200 | R&D Systems, #MAB2018 |
| TRA-1-81 | *PODXL* | mouse | 1:400 | Santa Cruz Biotechnology, #sc-21706 |
| ZEB2 | *ZEB2* | mouse | 1:50-1:100 | Santa Cruz Biotechnology, #sc-271984 |
